# Intersection of transient cell states with stable cell types in hippocampus

**DOI:** 10.1101/2025.10.17.683151

**Authors:** Jack A Olmstead, Lauren E King, Brenda L Bloodgood

**Affiliations:** Neurosciences Graduate Program; Medical Scientist Training Program; Department of Neurobiology, School of Biological Sciences; UC San Diego School of Medicine, La Jolla, CA, United States

## Abstract

The transcriptome of a brain cell encodes both its stable identity and its dynamic responses to environmental stimuli. While significant progress has been made in categorizing cell types within the brain, deciphering to what extent transcriptional identity and transcriptional state are related remains a major technical and conceptual challenge. Here, we present a single-nucleus RNA-sequencing atlas of the mouse hippocampus spanning physiological and pathological stimuli and multiple circadian phases, enabling unified analysis of activity-, circadian-, and cell-type-dependent transcriptional programs. Taxonomically assigned cell types are largely stable despite the induction of different activity states, with a notable exception in the dentate gyrus. Activity and circadian rhythm each drive robust, largely nonoverlapping transcriptional responses, with convergent regulation on genes involved in specific pathways, including endocannabinoid signaling, excitability, and chromatin remodeling. These results underscore the necessity of integrating cell-type taxonomy with transcriptional state to capture how diverse cell types respond to experience.

## Introduction

Activity-dependent gene expression is a fundamental mechanism that neural cell types use to respond to experience and coordinate adaptive cellular responses. For canonical immediate-early genes (IEGs) such as *Fos*, *Egr1*, and *Npas4*, transcription begins to rise within minutes and peaks within an hour, highlighting the transient kinetics of these programs^1–6^. Large-scale transcriptomic atlases have provided precise molecular definitions of cell types in the brain^7,8^, but interpretation of IEG expression within these foundational datasets has been fraught with known artifacts produced during cell capture, a process which itself induces IEGs^9,10^. A concerted effort is therefore necessary to characterize the intersection of activity states with a standardized taxonomy of cell types.

The hippocampus is a central structure for learning and episodic memory^11^, and displays strong cell-type-specific network activity patterns during exploration and rest^12–17^. Behavioral states that drive plasticity—such as exploration during arousal and sharp-wave ripples during rest or sleep—predominate at different points in the circadian cycle, positioning the hippocampus as a model system for developing conceptual approaches to the integration of transcriptomic cell types and multidimensional cell states. Prior bulk RNA-seq studies in mice have shown circadian cycling of clock genes in the hippocampus^18,19^, but no single-cell data exist to determine how neurons and glia engage these programs. Circadian phase influences excitability^20–23^, seizure susceptibility^24,25^, and memory performance^18,26^, making the interaction between activity-dependent and circadian-dependent transcription biologically important, yet poorly characterized. To date, transcriptional rhythms across the circadian cycle have not been described in the hippocampus using single-cell RNA-seq approaches, leaving open how different cell types entrain their gene expression to this important biological variable.

In this study, we leverage state-of-the-art cell-type resources and a multifactorial experimental design to create an atlas of hippocampal cell states across activity conditions and circadian timepoints, which we call the Activity and Circadian Transcriptomes Dependent on Physiological and Pathological activity (ACT-DEPP) dataset. (Note: Initial analysis of these data was published in a dissertation by J.A.O.^27^) With this resource, we show that enriched-environment (EE)- and seizure-induced neuronal activity engage qualitatively different transcriptomic programs within all cell types of the hippocampus. We find that within the Allen Brain Cell taxonomy of cell types, there is a collapse of cell type and activity state into a unitary identity among a subset of dentate granule cells. We describe a novel suite of marker genes that are transiently induced by activity in subpopulations of excitatory, inhibitory, and glial cells in the hippocampus. We report a gradient of IEG induction along the dorsal-ventral axis of CA1 in response to EE exposure. Finally, we show here that circadian rhythm coordinates widespread transcriptomic programs in both hippocampal neurons and glia at baseline. By comparing EE exposure to baseline states across the diurnal cycle, we systematically describe how activity- and circadian-dependent transcriptional programs intersect within hippocampal cell types and show that these variables influence endocannabinoid signaling, cell excitability, and chromatin remodeling.

By integrating transcriptional signatures of activity and circadian time into a taxonomy of cell types within the ACT-DEPP dataset, we provide both a conceptual framework for understanding state-dependent gene regulation and a resource for probing how activity patterns shape hippocampal plasticity across the circadian cycle.

## Results

### Generation of the ACT-DEPP dataset

Single-nucleus RNA sequencing (snRNA-seq) using SPLiT-seq^28^ was performed on hippocampi from animals exposed to a standard environment (SE); an enriched environment (EE) for 30 minutes or 6 hours; or kainic acid-induced seizures (KA) for 30 minutes or 6 hours. The resulting dataset includes 93,066 nuclei from 48 biological replicates, capturing gene expression in baseline, physiological, and pathological activity conditions across circadian timepoints (**Fig. 1A-B; Figure 1—figure supplement 1**).

**Figure 1.**
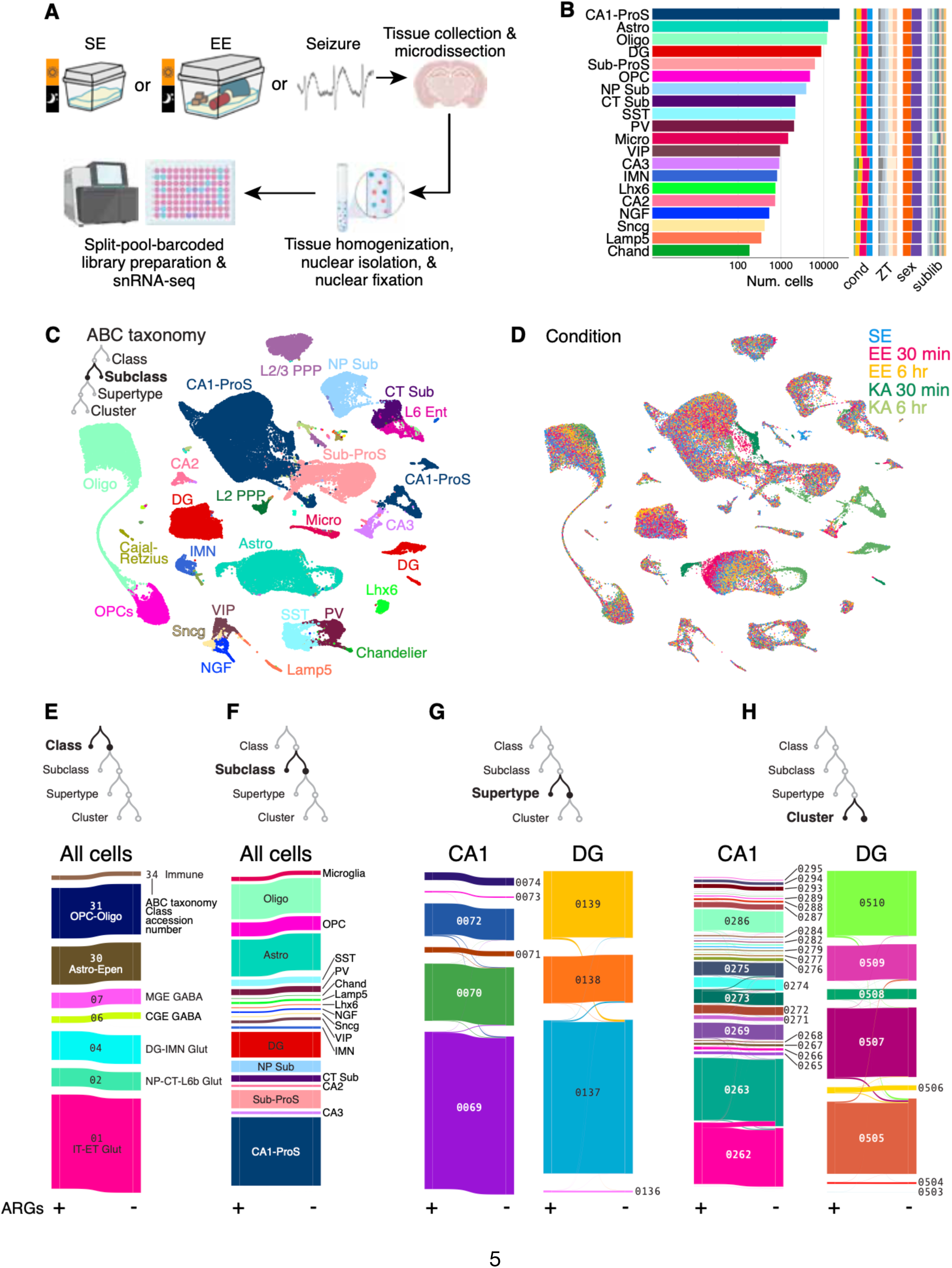
Generation of the ACT-DEPP dataset. (A) Experimental schematic of generation of the ACT-DEPP dataset. (B) Distribution of nuclei number, experimental conditions, and sample metadata among major subclasses. Some neocortical and subiculum Subclasses appearing in Figure 1D have been omitted from this panel, as they were not analyzed in any further downstream analyses. (C) UMAP embedding of ACT-DEPP nuclei colored by subclass. Inset: Single-nucleus transcriptomes were mapped to the Allen Brain Cell (ABC) whole-brain atlas, which assigns cells in a hierarchical taxonomy to one class, subclass, supertype, and cluster. (D) As in (C), but colored by activity condition. (E-H) Sankey flow diagrams showing changes in ABC-taxon identity for nuclei before (+) and after (-) removing ARGs from single-nucleus transcriptomes. (G) and (H) show the effect at lower ABC taxa for the two largest neuronal subclasses, CA1 and DG. *CA1-ProS, cornu ammonis 1-prosubiculum; L2/3 PPP, layer 2/3 pre-, pro-, parasubiculum; NP Sub, near-projecting subiculum; CT Sub, corticothalamic subiculum; L6 Ent, layer 6 entorhinal; DG, dentate gyrus granule cells; Sub-ProS, Subiculum-prosubiculum; Micro, microglia; Astro, astroglia; Oligo, oligodendrocyte; OPC, oligodendrocyte precursor cells; Epen, ependymal cell; NGF, neurogliaform cell; IMN, DG immature neuron; SST, GABAergic somatostatin-expressing interneuron; PV, GABAergic parvalbumin-expressing interneuron; GABAergic vasopressin internal peptide-expressing interneuron; Sncg, GABAergic Sncg-expressing interneuron; Lhx6, GABAergic Lhx6-expressing interneuron*

Quality-controlled nuclei were annotated using MapMyCells and integrated with the Allen Brain Cell (ABC) taxonomy^7^, assigning each nucleus to hierarchical ranks of Class > Subclass > Supertype > Cluster (**Fig. 1C inset; Figure 1—figure supplement 2-4**). Note: the Subclass rank generally corresponds to conventionally understood cell types within the hippocampus, e.g., dentate granule cells, CA1 pyramidal cells, PV-expressing basket cells, etc. The UMAP embedding recapitulated relations of canonical hippocampal cell populations at the Subclass level (**Fig. 1C**), with additional clear transcriptional distinctions for both KA and EE conditions within neuronal and glial Subclasses (**Fig. 1D**).

To confirm that cell-type annotation was independent of activity-induced transcriptional states, a set of known activity-regulated genes^2^ (ARGs) were removed, and annotations were remapped. Cell-type re-annotations at the Class and Subclass rank were highly stable (**Fig. 1E**-**F**), with minimal reassignment even at the lower ABC taxonomic ranks of Subclass and Cluster (**Fig. 1G-H**). The ACT-DEPP dataset thus robustly registers hippocampal cell types to the ABC taxonomy and reliably distinguishes transcriptional states driven by physiological and pathological stimuli.

### The activity-dependent transcriptome obscures cell identity in DG

Seizures have been used extensively to study ARG expression in the past, but direct comparisons to naturalistic stimuli are limited^29,30^. Moreover, the near-complete segregation of KA populations within some Subclasses in our uniform manifold approximation and projection (UMAP) plots led us to hypothesize that seizures constitute a categorically different transcriptional state from physiological manipulations. We therefore analyzed differentially expressed genes (DEGs) in CA1 and DG, the most abundant excitatory neuron Subclasses in our samples, at 30 minutes and 6 hours. KA induced an order of magnitude more DEGs than EE at both time points. Approximately one-third of EE-induced DEGs at 30 minutes were distinct from those induced by KA (**Fig. 2A**). At 6 hours, CA1 responses were fully shared between EE and KA, while DG maintained a unique DEG set specific to EE (**Fig. 2B**). Thus, novel experience and seizure induce transcriptional programs that differ greatly in magnitude and, apart from CA1 after 6 hours, contain stimulus-specific genes.

**Figure 2.**
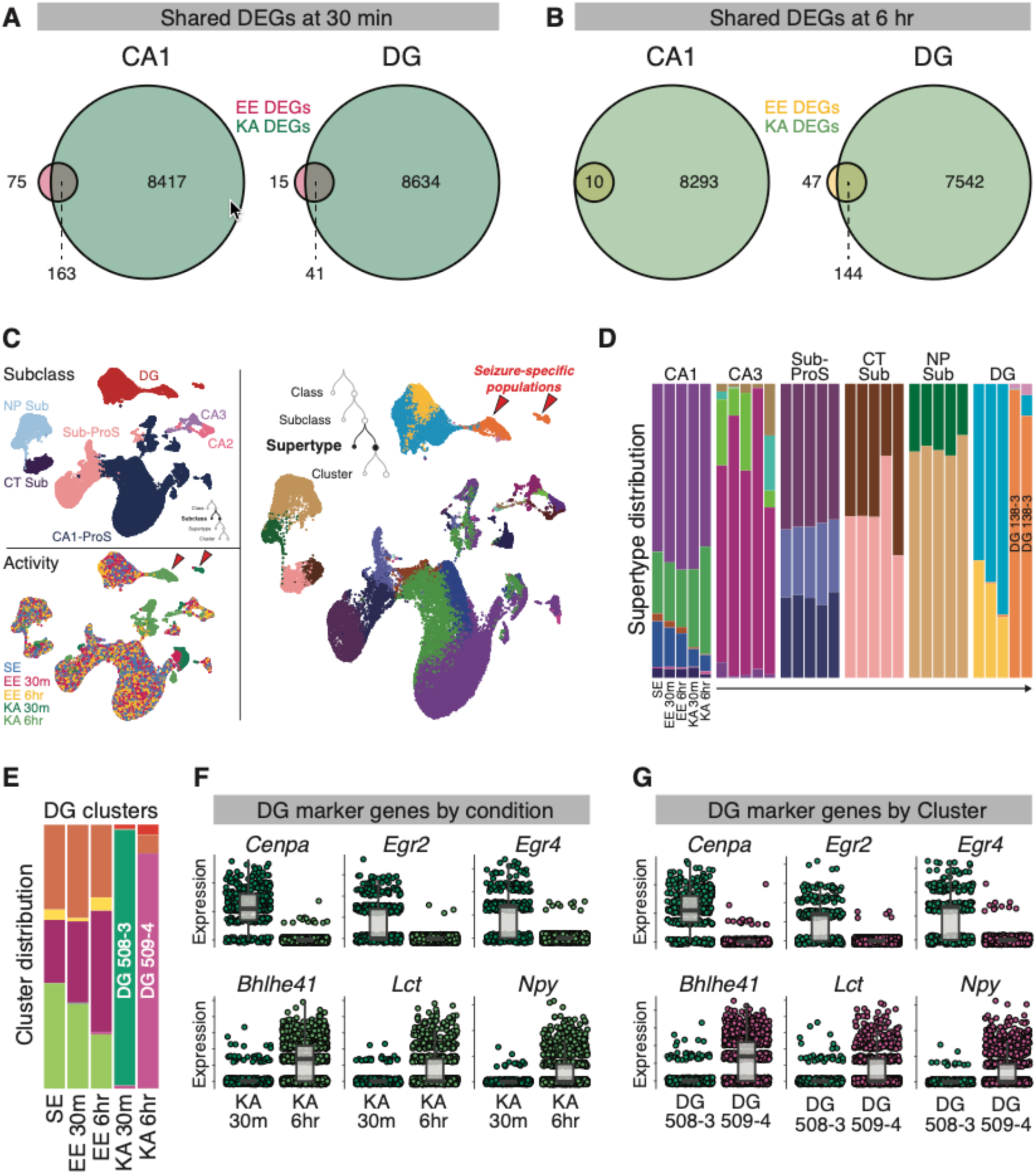
The activity-dependent transcriptome obscures cell identity in DG. (A-B) Venn diagrams of shared DEGs between KA or EE vs SE at 30 min (A) or 6 hr (B) in CA1 (n_EE30m_ = 3329 nuclei, n_SE_ = 1907 nuclei, n_KA30m_ = 354 nuclei) and DG (n_EE30m_ = 7119 nuclei, n_SE_ = 6754 nuclei, n_KA30m_ = 961 nuclei). DEGs were defined as genes with absolute log2(FC) > 1.5 and FDR < 0.05 using pseudobulk expression (Methods). (C) Re-embedded UMAPs of excitatory neuronal nuclei colored by Subclass (left, top), activity condition (left, bottom), or Supertype (right). Cross-reference of the UMAPs reveals KA-specific populations marked by red arrows. (D) Distribution of Supertypes within each subclass across different conditions. Colors within each column represent proportions of different Supertypes within the labeled Subclass.(E) As in (D), but for DG Clusters only. (F-G) Single-nucleus, SC-transformed expression (Hafemeister & Satija 2019) of marker genes for DG Cluster identity, plotted by condition (F) or Cluster (G)

To confirm that DEGs ascertained with DESeq2 were robust to various statistical frameworks, we also used limma-voom and fitted negative-binomial linear mixed models (NB-GLMMs) to normalize and estimate expression for DGE analysis. We first used limma-voom to revisit the DGE analysis in **Figure 2A-B** (**Figure 2—figure supplement 1**). Interestingly, there were no DEGs detected by limma-voom in the CA1 population after EE 6h; otherwise, results were qualitatively similar to results found using DESeq2. Next, we examined whether estimations of fold-change between DESeq2, limma-voom, and NB-GLMMs were quantitatively comparable. We observe that EE-induced changes in expression were comparable for canonical IEGs within CA1 and DG, albeit with more conservative IEG fold-change estimates when using DESeq2 shrinkage under certain conditions when compared with the other two methods (**Figure 2—figure supplements 2-3**). When comparing DGE methods genome-wide, concordance was high between gene lists ranked by activity-induced fold-change and statistical significance in the EE30m vs SE comparison (*ρ* = 0.985 in DESeq2 vs limma-voom, *ρ* = 0.985 for DESeq2 vs NB-GLMM; **Figure 2—figure supplement 4**), with a slight decrease in the EE6h vs SE comparison (*ρ* = 0.964 in DESeq2 vs limma-voom, *ρ* = 0.983 for DESeq2 vs NB-GLMM; **Figure 2—figure supplement 5**).

To look more closely at the effects of activity on different excitatory-neuron populations, we re-embedded UMAPs and color-coded nuclei by condition or Subclass/Supertype (**Fig. 2C**). Supertypes were clearly segregated within Subclass clusters, and strikingly, distinct Supertypes emerged as seizure-specific populations. This effect was particularly pronounced in DG, identified by its dramatically altered Supertype distributions between activity conditions (**Fig. 2D**). In DG, nearly all KA cells clustered within Supertype 138-3, with notable dissociation into Clusters 508-3 (KA 30m; green) and 509-4 (KA 6hr; purple, **Fig. 2E**). When we examined the spatial distribution of Supertype 138-3 within the ABC MERFISH atlas, we observe that this small population is distributed throughout the DG with a bias toward the dorsal hippocampus (**Figure 2—figure supplement 6**). Analysis of Cluster-defining genes revealed six marker genes strongly associated with seizure timing and cluster identity (**Fig. 2F-G**): *Egr2, Egr4, Cenpa, Bhlhe41, Lct, Npy*. Thus, Supertype DG 138-3 labels recently active dorsal granule cells, with additional discrimination of early and late transcriptional responses to activity at the Cluster level. This would likely not be detectable without both EE and KA conditions, given the sparsity of DG activation during naturalistic activity^31–35^.

### EE exposure induces DEGs within IMNs after 6 hours but not 30 minutes

EE exposure has been previously shown to induce postnatal neurogenesis within the dentate gyrus. Even though the current experimental timepoints were not designed to capture the emergence of adult-born granule cells caused by EE exposure, we were interested to see whether EE induced acute changes in gene expression within the preexisting immature neuronal (IMN) population. IMNs exhibit no DEGs following EE 30 min, but exhibit several following EE 6h (**Figure 2—figure supplement 7**). Several DEGs such as *Fmnl2, Lypd6, Fat4,* and *Sema3e* are involved in cytoskeletal organization or intercellular adhesion, while others such as *Chrna7, Gabbr2,* and *Kcnh7* encode neurotransmitter receptor subunits or transmembrane channels. Together, these results suggest that even 6 hours of EE exposure may be sufficient to induce gene expression that ultimately facilitates network incorporation of this nascent population.

### Discovery of cell-type-specific ERGs induced by novel experience

To understand how different cell populations respond to naturalistic activity, we compared early-response gene (ERG) expression across major groups of hippocampal Subclasses (DEGs were categorized as either ERGs or LRGs based on their peak- expression timing; see Methods and ref ^10^). Both excitatory and inhibitory neurons share broad upregulation of canonical ERGs such as *AP1*-, *Egr*-, *Nr4a-*, and *Dusp*-family genes, as well as *Arc*, and *Npas4* across nearly all Subclasses examined (**Fig. 3A, E**); these genes exhibit some of the strongest induction genome-wide across nearly all Subclasses, including astrocytes and microglia (**Fig. 3B, F, I**).

**Figure 3.**
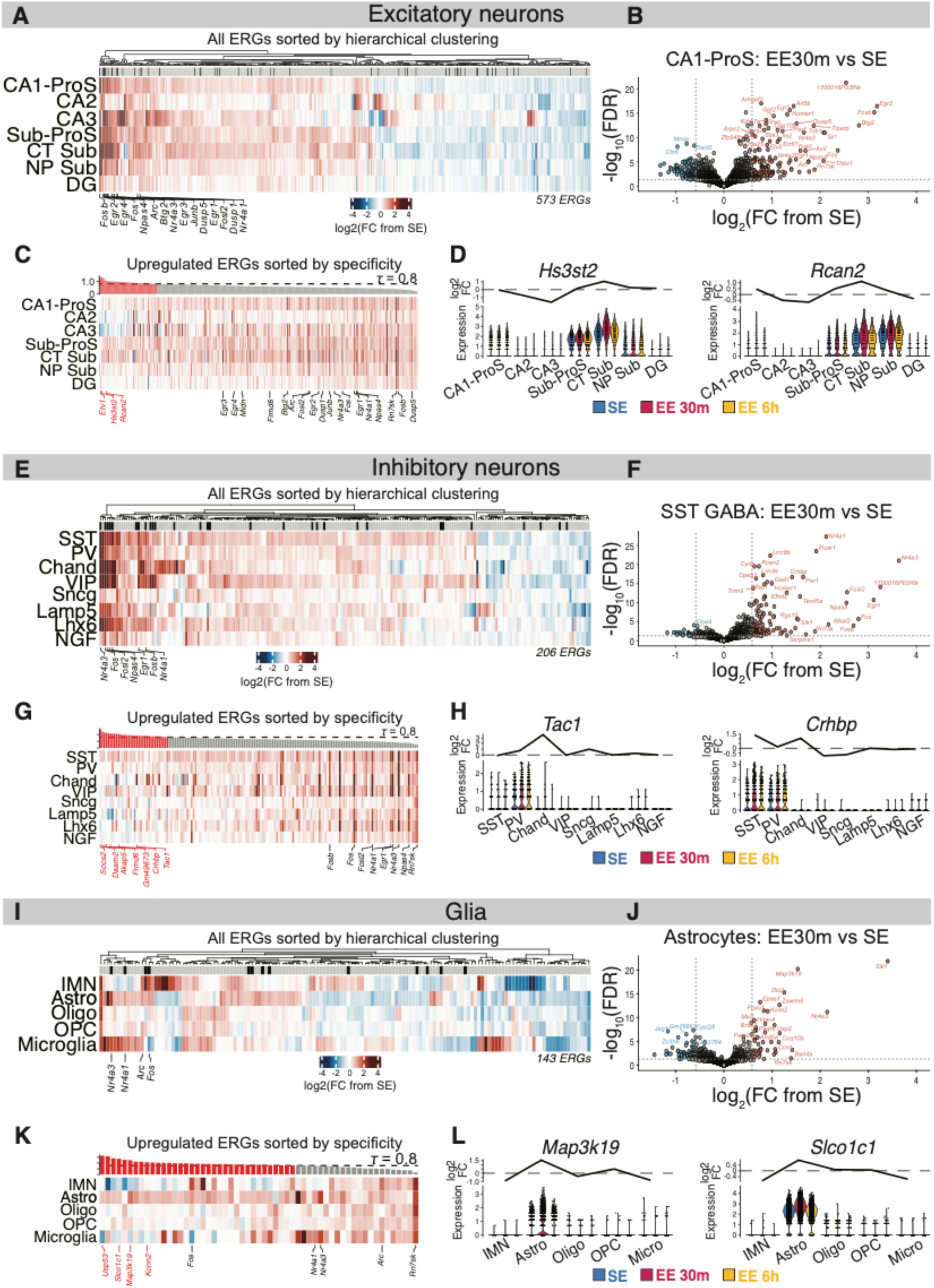
Discovery of cell-type-specific ERGs induced by novel experience. (A) Change in expression in EE30m vs SE for all ERGs in excitatory neuronal subclasses; gene columns are ordered by hierarchical clustering, and genes encoding transcription factors are annotated with black ticks above heatmaps. For every Subclass, we performed pseudobulk DEG analysis at EE 30 min vs SE and EE 6 hr vs SE. The heatmap contains all DEGs—ascertained from any Subclass listed—that were classified as ERGs. ERGs are defined as DEGs whose fold- change in EE vs SE was higher after 30 min instead of 6 hr (Methods). (B) Volcano plot of DEGs in CA1-ProS nuclei at EE30m (n = 3329 nuclei) vs SE (n = 1907 nuclei). (C) Heatmap of upregulated ERGs in excitatory neuronal Subclasses, ordered by Subclass- specific induction (tau score; Methods). Canonical ERGs are annotated in black text, while novel example Subclass-specific genes are annotated in red text.(D) Violin plots of SC-transformed expression for example genes with high tau scores. (E-H) as in (A-D) but for inhibitory neuronal subclasses. (I-K) as in (A-D) but for immature neurons and glial subclasses.

Next, we were interested in determining which upregulated ERGs exhibited specific activity-dependent induction in one or a few Subclasses. We calculated a tau specificity score previously developed for bulk RNA-seq count data^36^ to filter for cell-type specific changes in expression within EE 30 min vs SE across Subclasses (Methods). When doing so, we observe that canonical ERGs have some of the lowest cell-type specificity (**Fig. 3C, G, J**; **Figure 3—figure supplement 1**). Meanwhile, screening for high-tau ERGs reveals genes whose induction is specific to one or a few related cell types. For example, *Hs3st2* and *Rcan2* are both genes whose expression is (1) specific to spatially restricted subiculum Subclasses, (2) acutely upregulated following activity, and (3) decaying to baseline expression at 6 hours (**Fig. 3D**). Likewise, *Tac1* is an exemplar activity-dependent marker gene among PV-expressing GABAergic cells and while *Crhbp* is expressed in medial-ganglionic-eminence-derived interneurons broadly (**Fig. 3H**). Finally, we make special note of the astrocytes, surprising in their robust Subclass- specific response to acute EE exposure (**Fig. 3I-J**). Among the DEGs, *Map3k19* and *Slco1c1* stand out as genes highly specific to activated astrocytes (**Fig. 3K**). Thus, comparison of activity-dependent ERG expression following novel experience yields novel marker genes with highly cell-type-specific induction.

### Distribution of IEG expression across CA1-ProS Supertypes

Because CA1-ProS neurons were the largest Subclass in our dataset, we used the opportunity to explore experience-induced activity responses within sub-populations of these principal cells. First, we characterized activity in the entire population by adapting a previously used method^10^ to classify individual neurons as “active” or “inactive” based on thresholded induction of immediate early genes (**Fig. 4A**; **Figure 4—figure supplement 1**; Methods). In an independent implementation, we constructed a composite IEG score for each cell and observed that active/inactive classifications using this continuous score were highly concordant with classifications using thresholded IEG induction (**Figure 4—figure supplement 2**). Crucially, our library preparation captured only nuclear transcripts before they were exported to the cytoplasm, increasing the likelihood that active cells are classified as such specifically in response to the stimulus^3,9^. At baseline (SE), only 8% of cells were active. After 30 minutes of EE, activation increased to 33% of CA1 cells, then declined to 20% by 6 hours. KA seizure robustly activated nearly all cells at 30 minutes, decreasing to about half at 6 hours, with a bimodal distribution possibly reflecting ongoing versus resolved seizures^37^ (**Fig. 4B**).

**Figure 4.**
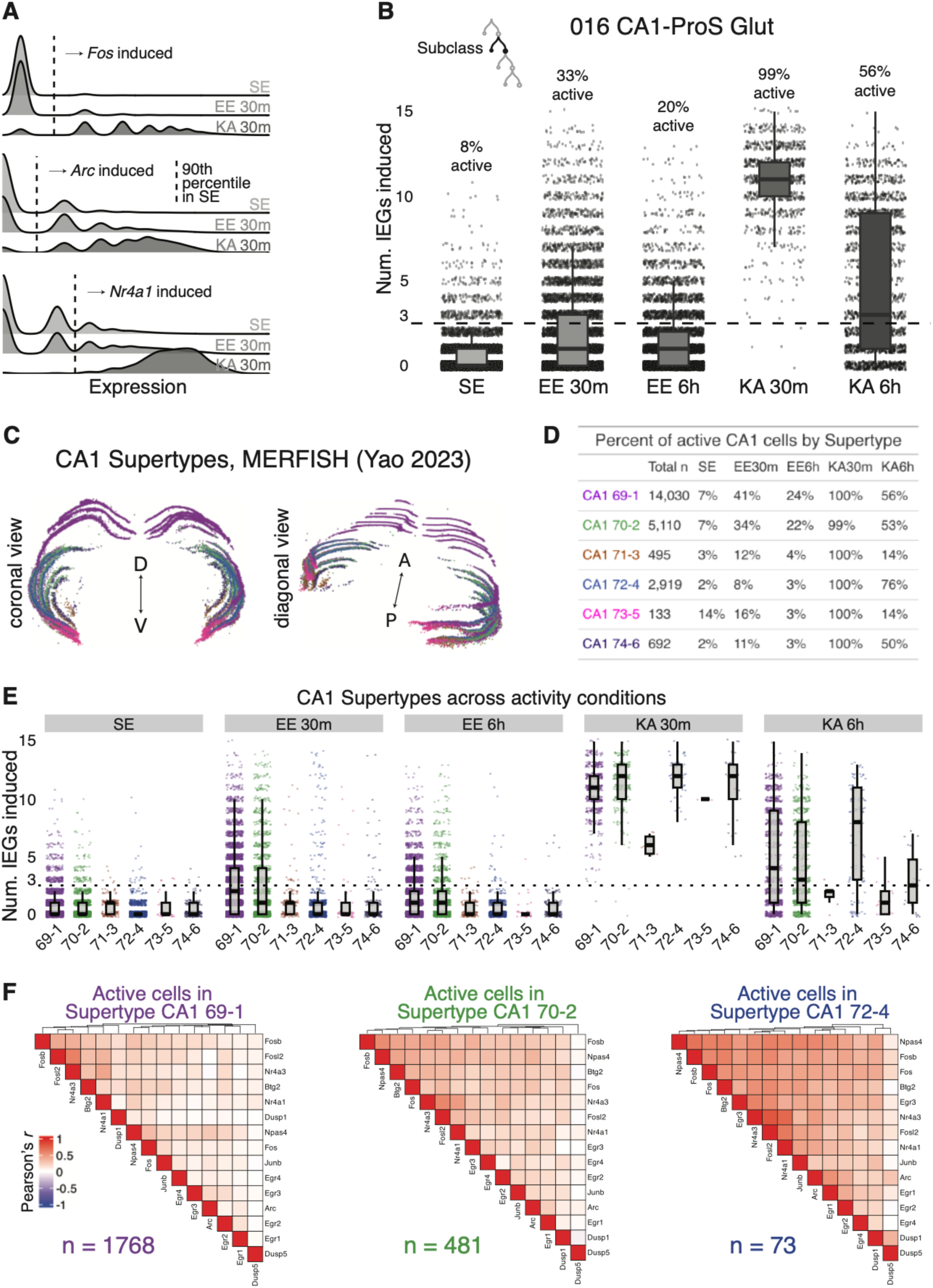
Distribution of IEG expression across CA1-ProS Supertypes. (A) Example plots demonstrating IEG induction thresholds using *Fos*, *Arc*, and *Nr4a1*. For a given gene, the induction threshold was defined as the 90th percentile of expression among SE nuclei; thresholds are represented as dotted lines.(B) Activation of single cells across activity conditions within CA1-ProS neurons. Active cells were defined as those that induced 3 or more IEGs, similar to the approach in Hrvatin et al. 2018 (Methods). (C) 3-D plots of ABC MERFISH brain slices (published data from Yao et al. 2023) with only CA1- ProS Supertypes plotted. (D-E) Activation percentages as in (B), de-aggregated by Supertype. (F) Gene-gene correlation matrices for IEG expression within active cells at EE 30 min. Dendrograms represent hierarchical clustering of genes.

Transcriptionally defined Supertypes correspond to gradations of anatomically distinct sub-populations of CA1 pyramidal neurons, evidenced by visualization within the ABC MERFISH atlas^7^ (**Fig. 4C**). We therefore hypothesized that we would observe differences between CA1 Supertypes that may underlie functional specialization within the hippocampus along anatomical axes^38,39^. We find that EE strongly activates Supertypes 69-1 and 70-2 compared to others—notable given the dorsal distribution of these Supertypes within hippocampus (**Fig. 4C**) and the engagement of this part of the hippocampus during exploration^40–42^. In contrast, KA seizure manipulations demonstrate that all Supertypes are competent at inducing IEGs following 30 minutes of global activity, although variation in active cells at 6 hours suggests differences in ongoing neural activity, ongoing transcription, or mRNA export among CA1 Supertypes (**Fig. 4D- E**).

Prior single-cell studies have analyzed IEG expression in different neuronal cell types, including in the hippocampus, to discern whether there are IEG “modules,” i.e. genes that are co-induced together^8,10,30^. No such analyses have revealed any interpretable structure of IEG activation. We sought to determine if the unprecedented cell-type resolution among ABC taxa in our dataset would reveal patterns of IEG expression in CA1 subpopulations that were aggregated in prior analyses. If a pattern to stimulus-dependent IEG induction existed, it would emerge in recently active cells, appearing as highly correlated expression modules among IEG subsets, with anti-correlation between subsets. When we tested this hypothesis among active cells in our most abundant Supertypes after 30 minutes of EE, we observed patterns of modest positive correlation among nearly all canonical IEGs across all Supertypes (**Fig. 4F**). We therefore conclude there is minimal evidence for structured or modular expression of IEGs within subsets of CA1-ProS neurons.

### Circadian-dependent transcription among major hippocampal cell types in SE

Biological rhythms entrained to the diurnal light cycle are ancient cellular processes driven by transcription-translation feedback loops that occur in many tissues, including the mouse hippocampus^23,43^. Yet, circadian cycling of transcription remains poorly understood outside the suprachiasmatic nucleus, with no single-cell studies characterizing the extent to which this occurs among different cell types in the hippocampus. We therefore sought to characterize transcriptional changes occurring at four Zeitgeber time points (ZTs) throughout the diurnal cycle under baseline conditions using our snRNA-seq approach, focusing on the four largest Subclasses to reduce noise. Canonical clock genes *Per1*, *Per2*, and *Cry2* peaked at ZT12 (start of the dark cycle), whereas *Clock* and *Bmal1* had opposite periodicity at ZT0 (**Fig. 5A**), aligning with previous hippocampal studies^18,19,23,44–48^.

**Figure 5.**
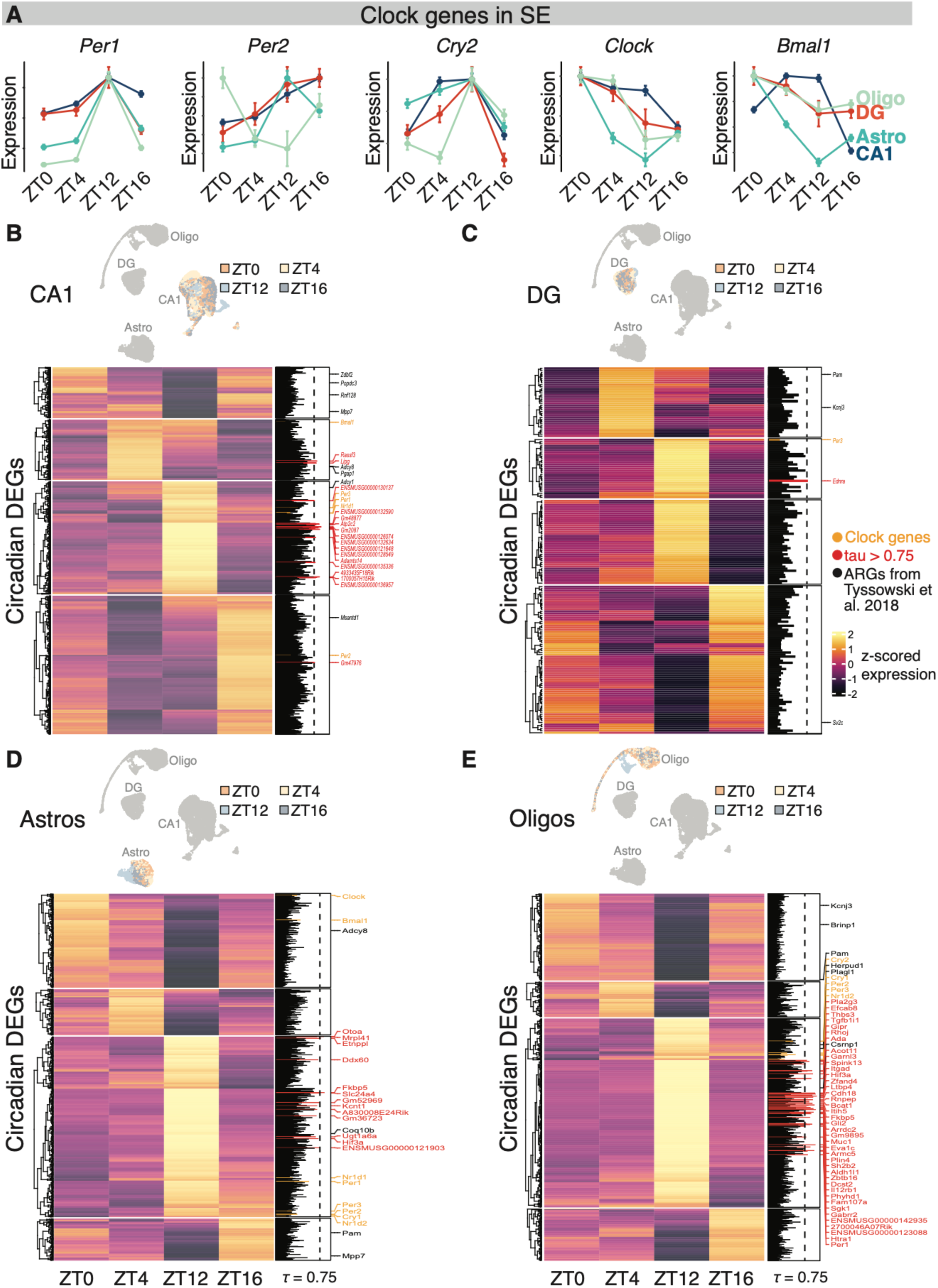
Circadian-dependent gene expression among major hippocampal cell types. (A) Pseudobulk expression of canonical clock genes across the circadian period in SE. Error bars represent s.e.m. (B-E) Above: UMAP embeddings of nuclei in circadian-DEG space, colored by ZT of collection (Methods). Below: Pseudobulk expression for circadian DEGs across the diurnal cycle. Clock genes (yellow), high-tau genes (red), and ARGs (black) are highlighted.

We next sought to characterize transcriptional changes taking place genome-wide using DGE analysis. When doing so, we capture the differential expression of canonical clock genes in every subclass tested (**Fig. 5B-E**, yellow annotations). We note again that *Per*- family expression peaks at ZT12 in every case but one. In addition to the *Per* family, its heterodimer partners *Cry1*, *Cry2*—as well as transcriptionally correlated Clock-gene regulators *Nr1d1*, and *Nr1d2*—peak at ZT12 in Subclasses where they are differentially expressed (**Fig. 5B-D**). Conversely, although we do not capture differential expression of *Clock* and *Bmal1* in every Subclass, their peak expression in such cases is not concurrent with *Per-*, *Cry-*, or *Nr1d*-family peaks. These phase-shifted expression peaks are functionally consistent with the canonical transcription-translation feedback loop that undergirds circadian rhythms^49^. Thus, the ACT-DEPP dataset captures circadian oscillations of transcription with high fidelity in multiple neuronal and glial cell types.

We next created UMAP embeddings of single-nucleus transcriptomes in circadian-DEG space (2725 genes across all Subclasses) to determine whether there is a transcriptomic structure tracking circadian time. We observe a clear separation of nuclei collected at ZT12 in every Subclass examined, with additional gradations of other timepoints in some cell types (**Fig. 5B-D**). High tau-specificity analysis confirms statistically what is visually apparent: In all Subclasses, ZT12 is a unique circadian-transcriptomic state with broad upregulation of many genes specifically at that time. Interestingly, in CA1-ProS cells, 12 of 17 of high-tau genes encoded lncRNAs, in contrast to DG granule cells (0/1), astrocytes (4/13) and oligodendrocytes (4/40). This points to a unique recruitment of the non-coding transcriptome in CA1-ProS neurons during this diurnal transition, the functional significance of which is unclear. (None of the 12 lncRNA gene symbols has any functional experimental data in the literature.) Finally, among previously characterized ARGs^2^ (ref ^50^ for *Adcy* genes), several of these varied throughout the circadian cycle, suggesting that there may be an interaction between ZT and activity within the hippocampus.

### Interactions between activity- and circadian-dependent transcription in the hippocampus

Previous studies have shown differences in canonical ARG expression at different times of day, although these studies could not make direct claims about stimulus-specific ARG induction due to their experimental design^18,18,19^. To directly test whether circadian time influences activity-dependent gene expression, we performed EE exposure at different ZTs. We hypothesized, based on these prior studies, that ARG induction would vary in at least some cell types across circadian time; that is, fold-change in ARG expression in EE vs SE would not be constant across the circadian period. We restricted our initial analyses to IEG and clock gene sets used in **Figures 4** and **5**, respectively. Surprisingly, there were no significant ZT:Activity interactions for IEGs in any of the Subclasses tested (**Fig. 6A-D**). This was unexpected, given previous reports of circadian-dependent IEG expression at baseline and upstream signaling cascades using non-RNA-seq methods^44,50,51^. We nevertheless note strong IEG induction in CA1, irrespective of ZT, unlike other Subclasses (**Figure 6—figure supplements 1-2**).

**Figure 6.**
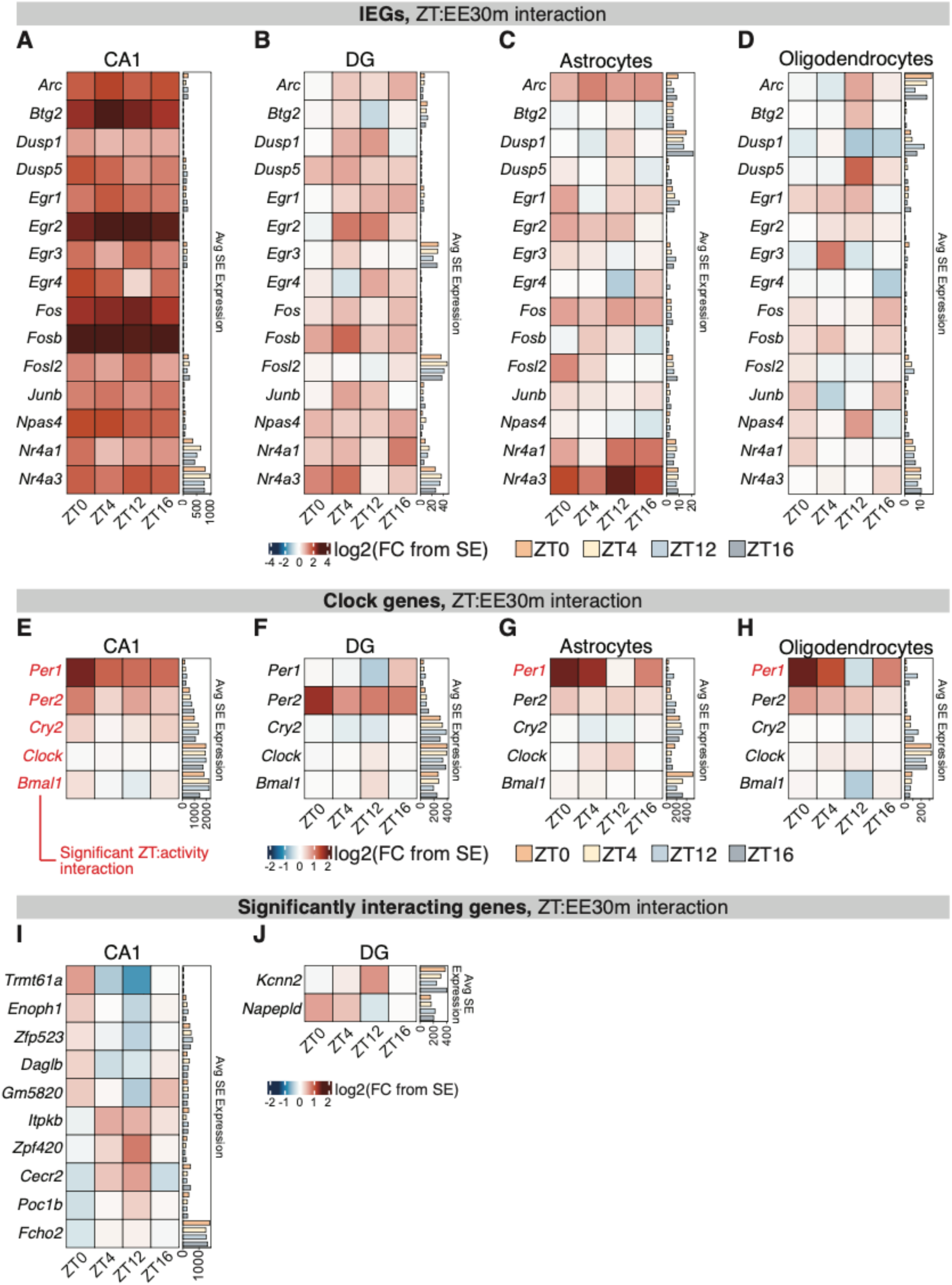
Interactions between activity- and circadian-dependent transcription in the hippocampus. (A-D) Expression change between EE30m vs SE for IEGs across the circadian period. Baseline pseudobulk expression for each gene at each ZT is plotted on the right.\ (E-H) As in (A-D) but for canonical clock genes. (I) Expression change between EE30m vs SE for genes with significant ZT:activity_condition interaction (Methods). Baseline pseudobulk expression for each gene at each ZT is plotted on the right. (J) As in (I) but for DG. Note: astrocytes and oligodendrocytes had no significantly interacting genes between ZT and activity condition.

Having established that the expression of clock genes varies substantially in baseline conditions (**Fig. 5A**), we hypothesized that, if present, activity-dependent induction of these genes would not be uniform across ZTs. Indeed, we find that not only are *Per1* and *Per2* inducible in response to EE exposure, but also that *Per1* exhibits a significant ZT:Activity interaction in all Subclasses except DG, with inducibility restricted to times excluding ZT12—notable because ZT12 corresponds to peak expression in SE (**Fig. 5A**). This observation can be interpreted as a “ceiling effect” of EE-induced expression mimicking the peak circadian expression.

Finally, we tested for ZT:Activity interaction genome-wide. Surprisingly, we only found a small number of genes whose expression was significantly modulated by ZT:Activity interaction, and these genes only appeared in the neuronal Subclasses (**Fig. 6I-J**). Among the cell types tested, we observe 13 genes that have a statistically significant interaction between ZT:EE exposure, none of which are canonical IEGs. Functional annotation of these genes reveals that two are involved in endocannabinoid synthesis (*Daglb, Napepld*), three in cell signaling or excitability (*Fcho2, Itpkb, Kcnn2*), and four in chromatin remodeling or gene expression (*Cecr2, Zfp420, Zpf523*). We therefore conclude that changes in gene expression following novel experience are relatively constant across the circadian cycle, although special mechanisms may be in place to selectively change induction of genes which modulate endocannabinoid synthesis and cell excitability within neurons of the hippocampus.

## Discussion

The ACT-DEPP dataset is an atlas of single-nucleus transcriptomes from the hippocampus, capturing transcriptional responses to both physiological and pathological brain activity. The creation of this atlas accomplishes two goals: (1) mapping of experience-dependent gene expression in a high-resolution cell-type taxonomy, and (2) characterization of circadian-dependent transcriptional oscillations in both neurons and glia of the hippocampus. All cell types are registered to a gold-standard brain-wide taxonomy, providing rich annotation that we leverage to demonstrate that stimulus-responsive cell states are embedded within broader cell-type identities, such as hippocampal sub-region (**Fig. 3**) and anatomical location (**Fig. 4**), with DG serving as an exception to this rule (**Fig. 2**). We identify the light-to-dark transition as a key inflection point for diurnal gene regulation in the hippocampus (**Fig. 5**) and show that activity-dependent induction does not vary widely across the circadian period (**Fig. 6**). We envision this resource as a roadmap for analyzing cell-type-specific, stimulus-responsive states in the brain using modern transcriptome-registration tools. More broadly, the ACT-DEPP dataset is a resource for understanding how physiological and pathological activity patterns drive experience-dependent transcriptional programs to support neural plasticity across the circadian cycle.

### Limitations

We take care to point out a limitation of our study: Dissociating neuronal gene expression driven by circadian phase from that driven by acute locomotion is inherently challenging because the two are tightly coupled in behavior and physiology. Locomotor activity co-occurs with changes in arousal, hormones, and neuromodulator release, many of which independently influence transcription, while the propensity for locomotor activity varies across the circadian cycle^23,52^. One argument that favors the distinction we make between activity- and circadian-dependent expression is that canonical IEGs do not overlap with circadian DEGs (**Fig. 5**), suggesting that activity programs and circadian programs may be induced by divergent regulatory mechanisms. Second, in contrast to neurons, glial transcription is less sensitive to rapid fluctuations in circuit activity and more influenced by systemic cues, making their rhythmic gene expression less confounded by locomotor events. Taken together, these observations support the validity of interpreting our findings as reflecting both activity- and circadian-driven transcriptional programs, rather than a single merged signal. Future studies which experimentally control for ambient illumination and/or locomotion—or at the very least measure the latter^52^ and include it as a covariate in RNA-count modeling—are required to dissociate contributions from locomotion and circadian rhythms on gene expression.

We also disclaim notable variance in cell number and read depth among a small number of our biological replicates (**Figure 1—supplementary figures 3-4**). Although our pseudobulk approach accounts for both of these factors by computing size factors for gene-count normalization before performing DGE analysis, it is nevertheless likely that estimates of variance are noisier in these conditions.

We relied on DESeq2 for all the DGE analysis in **Figures 1-6**. Pseudobulk DGE analysis has been shown to be the most robust against Type I errors for single-cell data while performing the best in balancing Type II error rate^53,54^. Thus, the results reported in this paper should be protected from unacceptably high false-positive rates. However, we do observe that DESeq2 shrinkage aggressively reduces estimated fold-change compared to limma-voom or NB-GLMM methods when replicate variance is high. For example, the DG population possesses unique biological properties, such as a smaller nucleus and sparser task-related activation compared to pyramidal cells, that contribute to technical sampling noise of low-abundance transcripts such as IEGs. The effect of resulting abundance measurements using single-cell sequencing is increased technical noise and likely over-shrinkage of fold-change estimations at the EE 6h timepoint (**Figure 2—figure supplement 2-3**). Thus, our findings should be interpreted as a conservative observation of activity- and circadian-dependent changes in gene expression.

### Cell states and cell types

Beyond our investigation of circadian transcription, our study has three main contributions to our understanding of activity-dependent gene expression. First, we extensively leverage registration of our dataset with the ABC atlas—a state-of-the-art ontology of cell types in the mouse brain^7^—to query activity-dependent states among high-resolution cell types. The ability to observe the effects of activity manipulations at the Supertype and Cluster level allows us to infer activity distributions along functionally relevant anatomical axes in the hippocampus (**Fig. 4**). Second, we distinguish several candidate genes that are highly specific not only to cell types, but also activity states (**Fig. 3**). These genes may serve as guideposts for the development of simpler next-generation tools to study activity-dependent processes with cell-type granularity going beyond IEG-based permanent labeling systems using *Fos, Arc, or Npas4*^55–58^—all of which are broadly upregulated across multiple cell types in our data, including non-neuronal cell types (**Fig. 3**). Such tools would have the advantage of being able to permanently access task-specific ensembles within specific cell types without the need for complex intersectional genetic strategies. Third, our descriptions of activity in baseline and “ceiling” conditions such as seizure provide strong evidence for the annotation of specific Supertypes and Clusters as highly temporally specific signatures of recent activity (**Fig. 2**).

Within DG granule cells, we observed a collapse of cell type and activity state at the supertype level (**Fig. 2**), and we also demonstrate that the Cluster rank in this population further resolves activity states into both acute and subacute activity histories, on the order of 30 minutes to 6 hours within our dataset (**Fig. 2E**). This example case in DG begs the question: Does the annotation of activity state—or any other condition-specific state, e.g. cellular manipulation or disease—at the Supertype or Cluster rank merit the collapse of these taxa with closely related inactive populations within the ABC taxonomy? More broadly, to what extent should reference atlases balance heterogeneity of cell states with the classification of cell types? On the one hand, the development of genetic tools to interrogate a particular Supertype or Cluster may produce unexpected results if that taxon represents a distinct cell state not present, or overrepresented, within an experimental dataset. On the other hand, it is easy to imagine how the annotation of active DG cells, for example, through a simple compositional analysis of Supertypes or Clusters would be useful for high-throughput readout of activity. It is our opinion that the correlation of specific cell states with lower ranks in the ABC atlas is acceptable, as long as these cell states are ascertained using robust experimental methods whose results are clearly disclosed to the community.

### Circadian rhythm changes baseline gene expression in the hippocampus

It has long been established that cells in the suprachiasmatic nucleus (SCN) use transcription-translation feedback loops entrained to ambient-light sensation to keep a “molecular clock” that tracks the diurnal cycle^59,60^. However, there are few investigations of hippocampal circadian-dependent transcription in the omics era, and no snRNA-seq studies that we are aware of. Therefore, we sought to characterize how transcription changes in the hippocampus across the circadian cycle using snRNA-seq. Our results extend previous work by confirming the presence of clock-gene cycling in hippocampal neurons and glia across the circadian cycle. The expression patterns of canonical clock genes we observe are consistent with earlier bulk hippocampal studies^18,19,23,44^, lending confidence that our sampling strategy captures genuine circadian processes rather than stochastic variation. Importantly, we find that ZT12 represents a distinct transcriptional inflection point across multiple cell types, coinciding with the transition from the light to dark period. This time is behaviorally salient for nocturnal animals such as mice, as locomotor activity begins to rise sharply before and during the onset of darkness^23,47,52,61^. Our data therefore support the interpretation that circadian gene expression in hippocampal cells is tightly coupled to this environmental and behavioral transition.

The similarity of transcriptomic profiles at ZT16 and ZT0 further suggests that circadian-driven transcription is concentrated within a narrow temporal window. Rather than a gradual progression across the cycle, many genes appear to undergo sharp, temporally restricted bursts of expression near the light–dark transition. This “bursty” model of circadian regulation is consistent with prior reports of pulse-like transcription in single cells^62–65^ and may reflect a strategy by which hippocampal cells track salient transitions without maintaining high levels of rhythmic transcription throughout the entire dark period.

Among the cell types tested, we observed a striking specificity in circadian-dependent transcription: oligodendrocytes, which exhibit relatively few activity-dependent DEGs (**Fig. 3**), display the largest number of circadian DEGs. This dissociation underscores that activity-dependent and circadian-dependent transcription represent partially non-overlapping regulatory axes. For oligodendrocytes, the high number of circadian DEGs suggests that diurnal regulation may play a dominant role in processes such as myelin turnover or sheath maintenance, at least during the developmental period tested (approximately P28 mice). In contrast, neuronal populations such as CA1 exhibit fewer circadian DEGs overall, with enrichment of lncRNAs at ZT12 compared to other cell types, hinting at separate mechanisms of regulatory control. Together, these results suggest that circadian transcription exerts heterogeneous but highly structured influence across hippocampal cell types.

### Circadian cycle has limited effect on activity-dependent transcription

We found no circadian differences in IEG induction (**Fig. 6A–D**), despite expecting higher expression in the dark period due to increased locomotor activity^52^, state-dependent hippocampal network events such as sharp-wave ripples during rest^14^, and prior reports of circadian oscillations in cAMP, MAPK, and CREB signaling^50,52,66,67^. Instead, IEGs were induced uniformly across the diurnal cycle in both neuronal and glial populations, exemplified by CA1 cells (**Figure 6—supplementary figures 1-2**). Prior work reported dark-phase upregulation of IEGs in rats under chronic EE^68^ or in cortex at ZT20^51^, but those studies differ in species, brain region, EE paradigm, stimulus, timing, and bulk vs. snRNA-seq, making direct comparison to our results inappropriate.

It has been previously shown that *Per* genes are inducible in the hippocampus in response to circuit activity^18,19,48^, a finding that we recapitulate after 30 minutes of EE. This finding, in conjunction with *Per*’s circadian oscillations in the absence of activity, raises important conceptual questions. What does it mean for there to be circadian-dependent expression when these genes are also induced by activity? We observe that at the highest peaks of EE-responsive induction, *Per* expression is comparable to its circadian peak. One speculation that could explain this phenomenon is that in the absence of activity, daily peaks in *Per* induction “guarantee” a dose of permissive regulatory-element access for PER-CRY target genes which would otherwise not be transcribed but are nevertheless necessary for cell function; activity-dependent *Per* expression would act as an adjuvant to predictable circadian bursts, possibly at genes important for cellular adaptations to circuit activity. Clearly, additional studies are necessary to understand the multifactorial causes and functional consequences of *Per* expression.

When testing for ZT:activity interaction genes, only 13 DEGs were found, all in neuronal subclasses. The functional classes of these DEGs suggest that there is a special role for fluctuations in intercellular endocannabinoid signaling throughout the circadian cycle, an enticing idea that may be relevant for neurological phenotypes. For example, despite the untold number of molecular etiologies for different epilepsies, circadian rhythms to seizure attacks are common in drug-resistant epilepsy, occurring in approximately 90% of patients^25^. This observation implies that there are circadian changes taking place in the brain that are not themselves pathogenic, but may affect seizure threshold across the diurnal cycle. In summary, our examination of the intersection of activity- and circadian-dependent gene expression is an important first step toward understanding the circadian transcriptome in the hippocampus, as it may canalize certain disease phenotypes.

## Author contributions

**Jack A Olmstead:** Conceptualization, Investigation, Methodology, Data curation, Formal analysis, Software, Visualization, Writing – original draft, Writing – review and editing

**Lauren E King:** Investigation, Writing – review and editing

**Brenda L Bloodgood:** Conceptualization, Supervision, Project administration, Funding acquisition, Writing – original draft, Writing – review and editing

## Supporting information

Supplemental Item 1

Supplemental Item 2

## Acknowledgements

We thank members of the Bloodgood lab and Dhanajay Bambah-Mukku for continuous discussion and feedback about experimental results and data analysis. We are indebted to Eric Halgren for providing computing resources. We thank Elizabeth Chamiec-Case and Lauren Valdez for their assistance with experimental troubleshooting. We kindly acknowledge the detailed experimental and analytic support provided by Brian Davis, Sam You, and Daniel Diaz of Parse Biosciences. Initial analysis of these data was published in a dissertation by J.A.O.^27^ This work was supported by funding awarded to BLB by the National Institute of Neurological Disorders and Stroke (R01 NS111162). This publication includes data generated at the UC San Diego IGM Genomics Center utilizing an Illumina NovaSeq X Plus that was purchased with funding from a National Institutes of Health SIG grant (#S10 OD026929).

The authors declare no competing interests.

## Materials and Methods

### Animals

Animal experiments were approved by the University of California San Diego Institutional Animal Care and Use Committee and followed guidelines according to the National Institutes of Health’s *Guide for the Care and Use of Laboratory Animals.* All experiments used wild-type C57BL6/J mice from The Jackson Laboratory (JAX#000664). Mice were approximately four weeks old, with balanced proportions of males and females distributed across control and experimental groups (**Figure 1—figure supplement 1**).

### Experimental procedures

#### Standard environment and circadian timepoints

Mice were group-housed in a vivarium on a 12-hour light/dark cycle and provided food *ad libitum*. To avoid non-experimental stimulation artifacts resulting from cage handling or transport, mice were transported from the vivarium to the standard environment (SE) at least 18 hours before experiments. The SE consisted of a plywood box paneled with sound-dampening foam, air-circulation fans, and LED lights whose power was controlled by a timer mimicking the vivarium’s light/dark cycle. Circadian timepoints were measured and reported on the 24-hour Zeitgeber time scale, with ZT0 marked as “lights-on” (at 06:00 PST) and ZT12 as “lights-off” (at 18:00 PST).

#### Enriched environment

For enriched-environment (EE) experimental conditions, animals were transferred from their home cage in the standard environment to a larger cage in a different room containing plastic and metal toys, tunnels, vertical scaffolds, and a running wheel. Mice were placed in the EE with at least one cagemate and left to explore continuously for the durations reported (i.e., for 30 minutes or 6 hours).

#### Kainic-acid seizures

Pharmacological seizures were induced using peritoneal injection of 15 mg/kg of kainic acid (KA; Santa Cruz, #SC-200454) as described previously^37,69^. Seizure onset was measured as the onset of rearing and accompanying bilateral forelimb clonus, approximately corresponding to Racine stage 4^70^. Mice were sacrificed either 30 minutes or 6 hours after seizure onset.

#### snRNA-seq sample collection

For sacrifice and sample collection, mice were deeply and rapidly anesthetized with isoflurane in an induction chamber. Following induction, the brain was removed from the skull into a dissection dish containing ice-cold dPBS (Gibco Cat. No. 14040133). Hippocampus was rapidly extracted from one hemisphere, and the approximate CA1 area was microdissected using a microblade. Although the purpose of this microdissection was to de-enrich samples of dentate granule cells, we still note that these cells nevertheless comprise the 4th largest subclass in our dataset (**Fig. 1B**). The dissected tissue sample was then placed in a microtube and flash-frozen. The other hemisphere was placed in a cryomold as a whole, embedded in OCT compound, and rapidly flash-frozen in isopentane chilled with dry ice. Left and right hemispheres were alternated for the collection of CA1 and whole-hemisphere collection, across sexes and conditions, to avoid any effects of lateralization (**Figure 1—figure supplement 1**). Both samples from each animal were stored at −70°C until use.

#### Nucleus isolation and fixation

Flash-frozen hippocampal dissectates were stored at −70°C until all experimental samples had been collected. Before nucleus isolation, all buffers, tubes, and instruments were pre-cooled on ice for 15-20 minutes before tissue was removed from −70°C. Reusable instruments such as the Dounce tubes and pestles (Millipore cat. #D8938- 1SET) were rinsed with RNAse-free PBS and then with RNAse Away detergent before being chilled on ice. All pipette tips (filtered tips only) and disposable tubes contacting samples were RNAse-free. All buffers used during nucleus isolation contained recombinant RNAsin to preserve RNA integrity (Promega, cat. #N2111), and actinomycin D to prevent artifactual transcription stimulated by the isolation protocol (Tocris, cat. #1229). Two main buffers were used for nucleus isolation: nucleus-isolation medium (NIM) and NIM-Douce-pestle buffer. NIM buffer consisted of 250 mM sucrose, 25 mM KCl, 5 mM MgCl2, 10 mM Tris-HCl (pH 7.4), 1 mM DTT, 0.2 U/uL RNAsin, 25 uM actinomycin D, and 1 cOmplete protease inhibitor mini tablet (Roche cat. #11836170001), dissolved in 10 mL of buffer. NIM-DP buffer consisted of 0.1% triton-X 100 detergent in NIM buffer.

To begin nucleus isolation, tissue samples were removed from −70°C and placed directly on ice to thaw. No more than five samples underwent isolation at any one time in order to minimize sample-handling time before fixation. First, 1 mL of NIM-DP buffer was added to the sample storage microtube, and the sample was transferred to a 2 mL Dounce tube using a 1000 uL pipette tip. Samples were homogenized using a Douce pestle in 1 mL of NIM-DP buffer, first with 10 strokes of the loose pestle, and then 17 strokes of the tight pestle. Two sets of dounce tubes and pestles were used at a time, and each was washed five times with PBS in-between samples. Dounced tissue homogenates were then spun in a fixed-angle benchtop centrifuge at 800 x g for 8 minutes at 4°C. The supernatant was removed, and the nuclear pellet, clearly visible, was resuspended thoroughly in 500 uL of NIM buffer using 30 strokes with a P200 pipette tip. Suspensions were then filtered through a 70 um filter before being spun at 200 x g for 10 min at 4°C in a swinging-bucket centrifuge. The supernatant was removed, and nuclei pellets were resuspended in 187.5 uL of Parse Biosciences’s “Resuspension Buffer” and strained through a 20 um filter to begin Parse’s Evercode Nuclei Fixation protocol (version 1.1, kit chemistry v3). In total, samples/nuclei were suspended in actinomycin D and kept on ice or in the pre-chilled centrifuge for the entirety of the isolation protocol, which took approximately 1 hour before proceeding directly to the Parse fixation protocol.

#### snRNA-seq barcoding and library construction

snRNA-seq library construction was performed using Parse Biosciences’s Whole Transcriptome v3 kit according to the manufacturer’s protocol. Briefly, nuclei originating from different samples are multiplexed and barcoded according to the SPLiT-pool combinatorial barcoding method^28^, wherein individual nuclei go through serial rounds of PCR barcoding, pooling, and splitting amongst a 96-well plate containing well-specific oligo barcodes. Split-pooling occurs in four serial barcoding rounds to ensure that each nucleus has a unique combination of barcodes ligated. All samples were loaded and barcoded together to avoid batch effects from separate library preparations. Libraries were sequenced at the UCSD Institute for Genomic Medicine on an Illumina NovaSeq X Plus using paired-end 100-bp reads. Sequencing was targeted at approximately 90,000 reads per nucleus, with a final average read depth of 66,702 reads per nucleus.

### Quantification and statistical analysis

#### Data preprocessing and quality control

Nuclei quality control was performed as outlined in **Figure 1—figure supplement 2**. Raw nucleus x gene count matrices from demultiplexed samples were generated from the FASTQ files using Parse Bioscience’s Trailmaker software pipeline, version 1.4.1. A “raw” Seurat object created by Trailmaker containing 110,728 nuclei, along with the digital gene-count matrix, associated sample metadata, and DoubletFinder scores^71^ was downloaded and used to begin nuclei QC. All following analyses were performed using R, version 4.4.1. Percent of reads mapping to mitochondrial reads was calculated for each nucleus. Nuclei that did not meet the following criteria were removed: transcript count (UMI) >= 1500; gene count >= 1500; and percent-mitochondrial-read count <= 5%. After this filtering step, 108,748 nuclei remained. Allen Brain Cell (ABC) taxonomic labels were assigned using the MapMyCells online tool. Next, single-nucleus raw gene counts were transformed using Seurat’s SCTransform function (v0.4.1), with the vars.to.regress argument set as the percent of reads mapping to mitochondrial genes (percent.mt) and SPLiT-Seq sublibrary. SCTransformation was computed across nuclei from all samples. (Note: Across-sample SCT-normalization, rather than within-sample normalization followed by integration was determined appropriate after normalizing, clustering, and visualizing UMAPs on a pilot sequencing run from this dataset, where we observed no batch effects due to sample or SPLiT-seq sublibrary). SCT-normalized counts were then used to perform principal component analysis (PCA), nearest-neighbor graph construction, initial Leiden cluster assignment, and uniform manifold approximation and projection (UMAP) embedding. This resulted in the segmentation of 108,748 nuclei into 47 Leiden clusters.

At this step, manual curation of 47 detected Leiden clusters was performed by experimenter J.O. cross-referencing plots and tables of UMI count, gene count, percent mitochondrial reads, doublet scores, and putative ABC cell type within each Leiden cluster. Based on these judgements, nuclei in 6 small Leiden clusters were removed because of low counts, high mitochondrial reads, high doublet scores, or the collision of many different putative ABC cell types. Finally, any nuclei with a DoubletFinder score < 0.02 were removed. After these quality-control steps, 93,066 nuclei remained (**Figure 1—figure supplement 2**). These again went through final PCA, Leiden clustering, and UMAP embedding to create the UMAP in **Figure 1** and final Seurat object (v5.2.0) uploaded to GEO.

#### ABC taxon assignment stability

Stability of ABC taxonomy assignment for single nuclei was performed by comparing two separate runs of the curated data through the ABC atlas MapMyCells tool. The first run was performed using the native mouse transcriptome. The second run was performed after removing ARGs that had been ascertained in a published study of activity-dependent gene expression^2^. The stability of assignment was then determined by comparing taxon equality within the same barcode and visualized with Sankey flow diagrams (**Fig. 1E-H**).

#### Differential gene expression

To identify differentially expressed genes (DEGs) across activity conditions, we performed pseudobulk DGE analysis independently for each Subclass using DESeq2^72^ for all main figures. For each Subclass, nuclei were subset and grouped by their sample of origin. Pseudobulk counts were aggregated at the sample level using Seurat’s AggregateExpression() function. Genes expressed in fewer than 1% of nuclei were excluded. DESeq2 was then used to construct a DESeqDataSet with sample-level counts and a design formula including activity condition (`expression ∼ activity_condition`). Only condition contrasts meeting a minimum per-group nucleus count threshold (n = 30 nuclei per condition) were retained for testing. Wald tests were performed, and log2 fold changes were shrunken using the apeglm method. Genes with absolute log2 fold change > 0.585 (corresponding to 1.5-fold change) and fdr < 0.05 were classified as differentially expressed. For analysis discovering DEGs between ZTs, log2 fold change values were not shrunken. This was done for ZT analyses but not activity analyses to minimize type II error rate; there are fewer biological replicates at each ZT (**Supplementary Table 1**). Results were saved separately for each Subclass and contrast.

In addition to DESeq2, we employed limma-voom (R package v3.60.6) and a negative-binomial generalized linear mixed model (NB-GLMM) to estimate expression for confirmatory DEG detection. Between-method estimated fold-change and statistical significance was compared between methods using forest plots for gene subsets of interest (**Figure 2—figure supplements 3-4**) and rank-rank hypergeometric overlap (**Figure 2—figure supplement 4-5**)^73^. Limma-voom was used to estimate pseudobulk expression used the formula

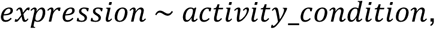

whereas NB-GLMM used the formula

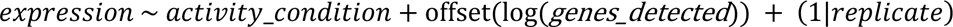

to protect against pseudoreplication bias using this single-cell DEG method.

#### Hierarchical clustering

For heatmap plots accompanied by a dendrogram, genes were ordered according to hierarchical clustering of the Euclidean gene distances using the package ComplexHeatmap^74^. In the heatmaps of **Figure 5**, genes were first clustered into 4 k-means clusters, and only after were organized according to hierarchical methods within each k-means cluster.

#### ZT:Activity interaction testing

To test for statistical interactions between the effect of ZT and the effect of activity on gene expression, we used the formula

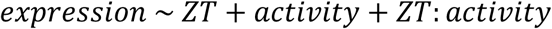

to test for a significant ZT:Activity interaction term with DESeq2. Pseudobulk gene counts were computed for each subclass separately. log2 fold changes were not shrunken in this analysis in order to increase sensitivity of this test. For *a priori* gene sets, multiple-hypothesis correction was performed by calculating the FDR among the *a priori* gene set. For example, the FDR rates reported in **Figure 6A-D** were corrected for 15 multiple comparisons for the set of 15 IEGs. Only genes with an FDR < 0.05 were labeled as significant DEGs.

#### ERG vs LRG classification

We used a similar method to a prior single-cell study to classify DEGs as mutually exclusive early-response genes (ERGs) or late-response genes (LRGs)^10^. Classification was performed separately for excitatory neurons, inhibitory neurons, and glial Subclasses. First, DEGs for all Subclasses and contrasts (EE30m vs SE, EE6h vs SE) were aggregated. Genes were provisionally labeled as ERG, LRG, or both based on differentially expressed in EE 30 min or EE 6 hr. Genes initially labeled as “both” were reclassified based on their peak fold-change timepoint. In cases where gene classification differed across Subclasses, the modal classification was assigned. Final classifications were saved for downstream analysis (**Figure 3**).

#### Tau calculation

We calculated a tau score for each gene, adapted from ref. ^36^, to discover and quantify highly cell-type specific gene expression. For example, a gene with a tau score of 0 would have promiscuous expression among the cell types examined, whereas a gene with a tau score approaching or greater than 1 would have highly cell-type specific expression. For analyses on non-fold-change expression data among different cell types (e.g. **Figure 5**), we directly applied the tau formula, in which tau was bounded by (0, 1):

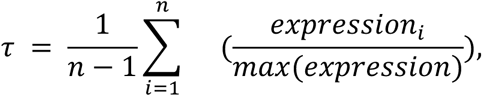

where *n* is the number of subclasses, and *expression_i_* represents the expression of the gene in subclass *i*; in these analyses, our tau threshold was set at 0.75 due to the index being bounded by (0, 1). For analyses on fold-change expression where expression values can be negative, the tau score was bounded by (0, ∞); for these analyses, our tau threshold was set higher at 0.8. In practice, tau scores above 0.75 were observed to be highly specific to one or a small number of subclasses.

#### Defining active cells

In order to determine the number of cells activated within a given condition, it was determined whether cells expressed the IEGs above a certain threshold: The IEG list was defined *a priori* based on the following 15 canonical IEGs: *Arc, Btg2, Dusp1, Dusp5, Egr1, Egr2, Egr3, Egr4, Fos, Fosb, Fosl2, Junb, Npas4, Nr4a1,* and *Nr4a3*. The induction threshold was determined for each gene individually, and was defined as the 90th percentile of expression in SE cells of the CA1 subclass. Thus, for each gene in the gene list, a binary classification was assigned as either induced or not-induced based on whether or not it surpassed the expression threshold (**Fig. 4A**). Finally, a cell was defined as “active” if it induced at least 20% of the genes (i.e., 3 or more) in the activity-regulated-gene list. To ensure that active-cell assignment is robust to minor changes in threshold parameters, we performed a parameter sweep varying gene number and expression threshold required for classification of an active cell. Active-cell fractions within CA1 were robust to minor changes in these thresholds and varied on a continuous gradient as expected (**Figure 4—figure supplement 1**).

A continuous IEG score was also used to classify active cells (**Figure 4—figure supplement 2**). The scale was constructed by (1) z-scoring a cell’s expression for a given IEG from the IEG’s mean/variance calculated in the SE then (2) summing the scaled expression of all 15 IEGs into a single score. Scaling was necessary to control for differences in transcript abundance among different IEGs (**Figure 4A**). Classifications determined using the continuous scale were compared to ground-truth classifications via the thresholding method used in **Figure 4**. ROC (R package pROC, v1.19.0.1)^75^ and PR curves (PRROC, v1.4)^76^ were constructed to visualize overall accuracy.

## Data and code availability

Fastq files, metadata for biological replicates, and raw and final Seurat files were deposited in GEO for public download (accession number GSE335423). All the code and package versions to reproduce this manuscript are publicly available at https://github.com/Bloodgood-Lab/Olmstead2026.

## FIGURE SUPPLEMENTS

**Figure 1 — figure supplement 1.**
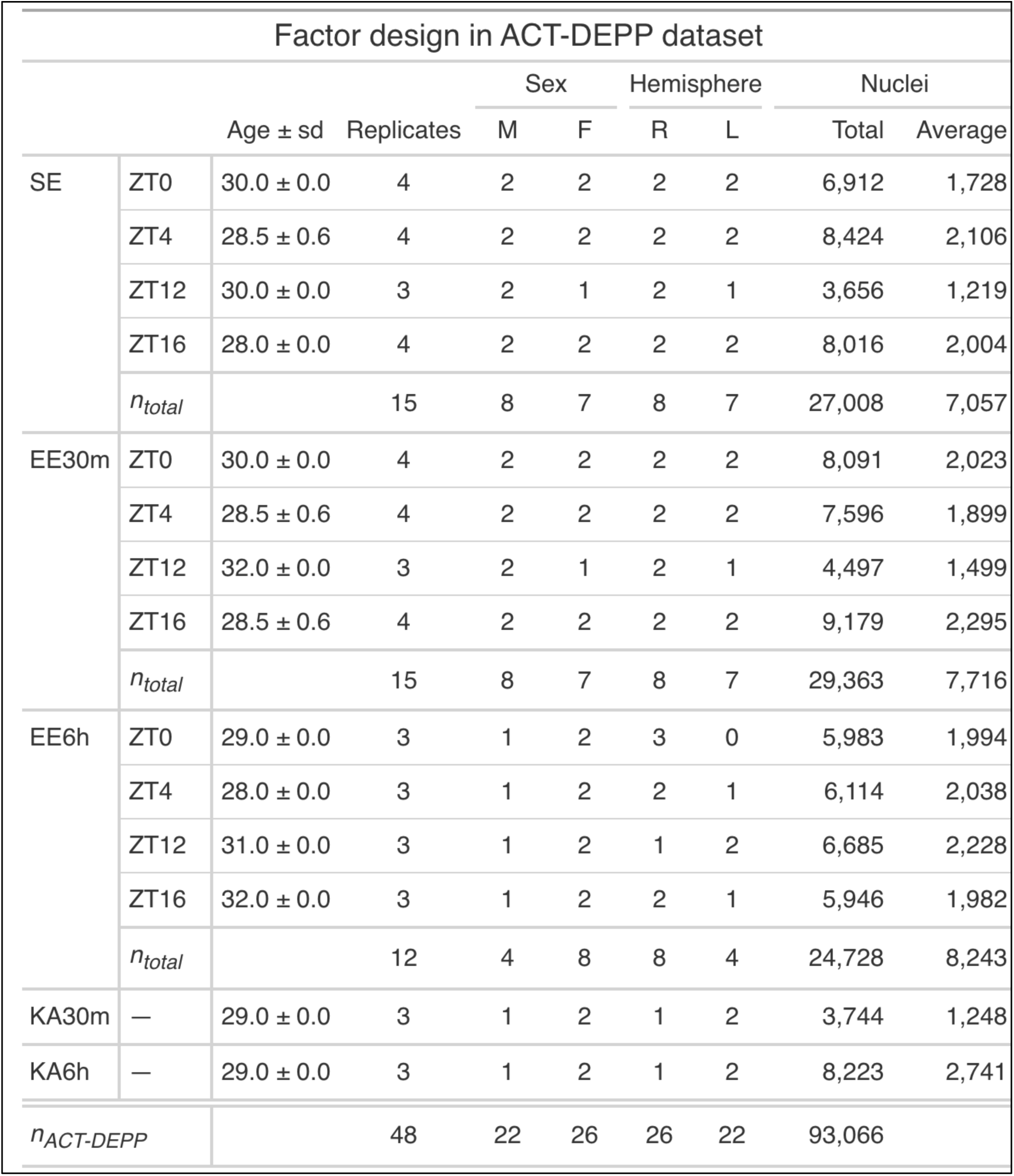
Experimental design and final nuclei yield among biological replicates. See also **Supplemental Item 1** for nuclei yields within major cell types.

**Figure 1 — figure supplement 2.**
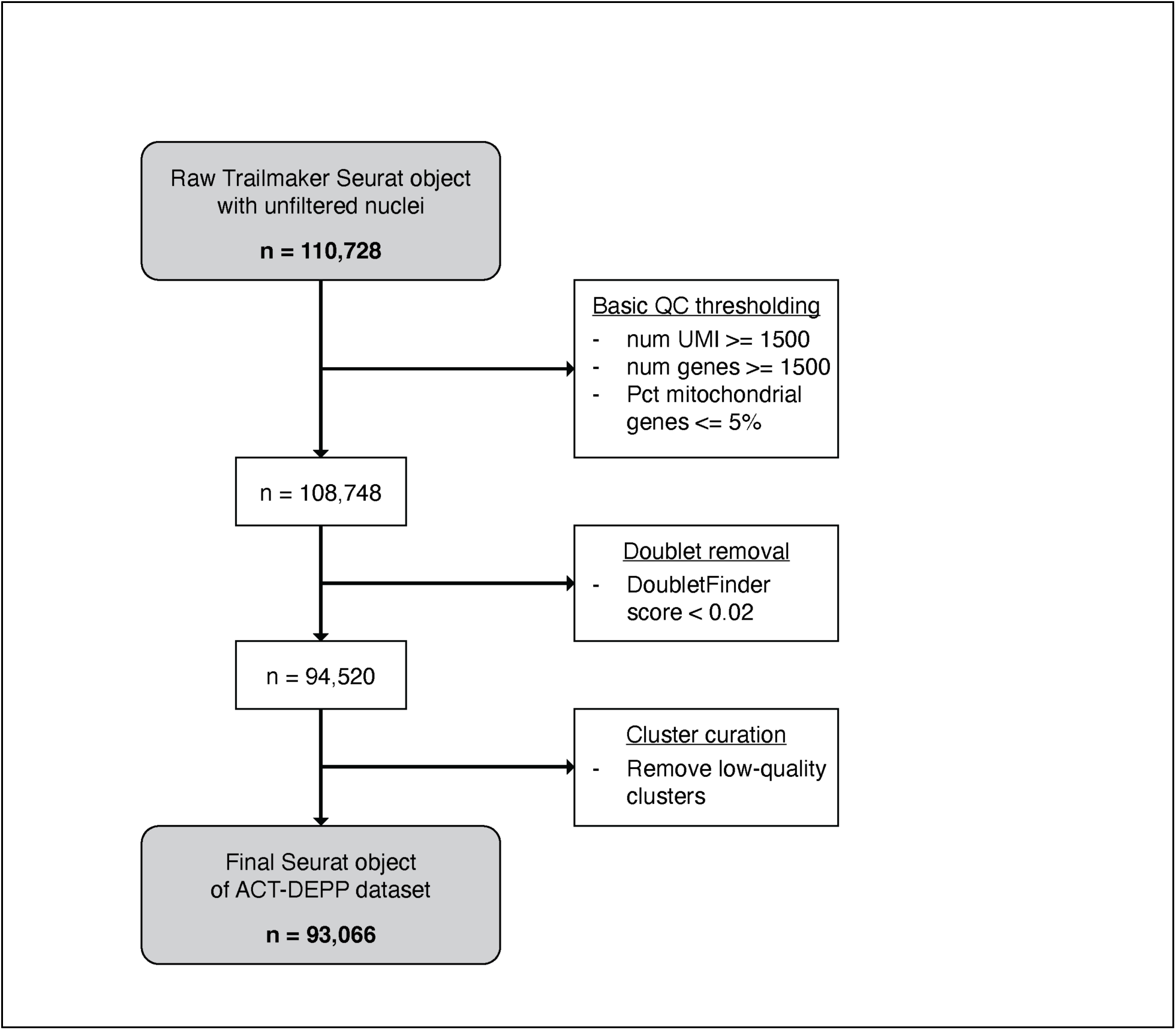
snRNA-seq quality-control workflow. Parse Biosciences’s Trailmaker version 1.4.1 was used to align barcoded reads and assemble raw nucleus x gene-count matricies. Intermediate downstream analyses were performed in Seurat v5.2.0.

**Figure 1 — figure supplement 3.**
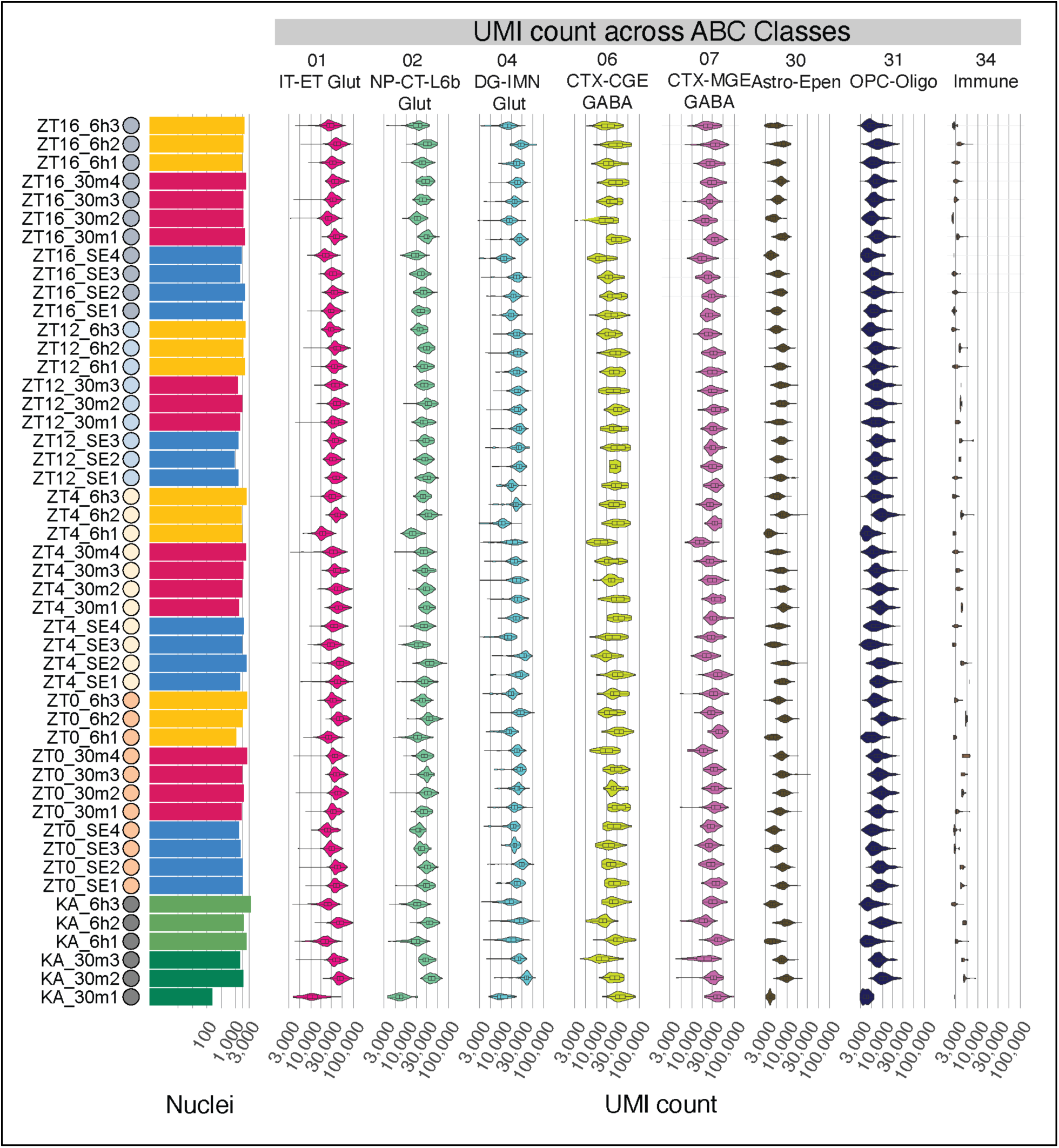
UMI counts among ABC cell Classes for each biological replicate.

**Figure 1 — figure supplement 4.**
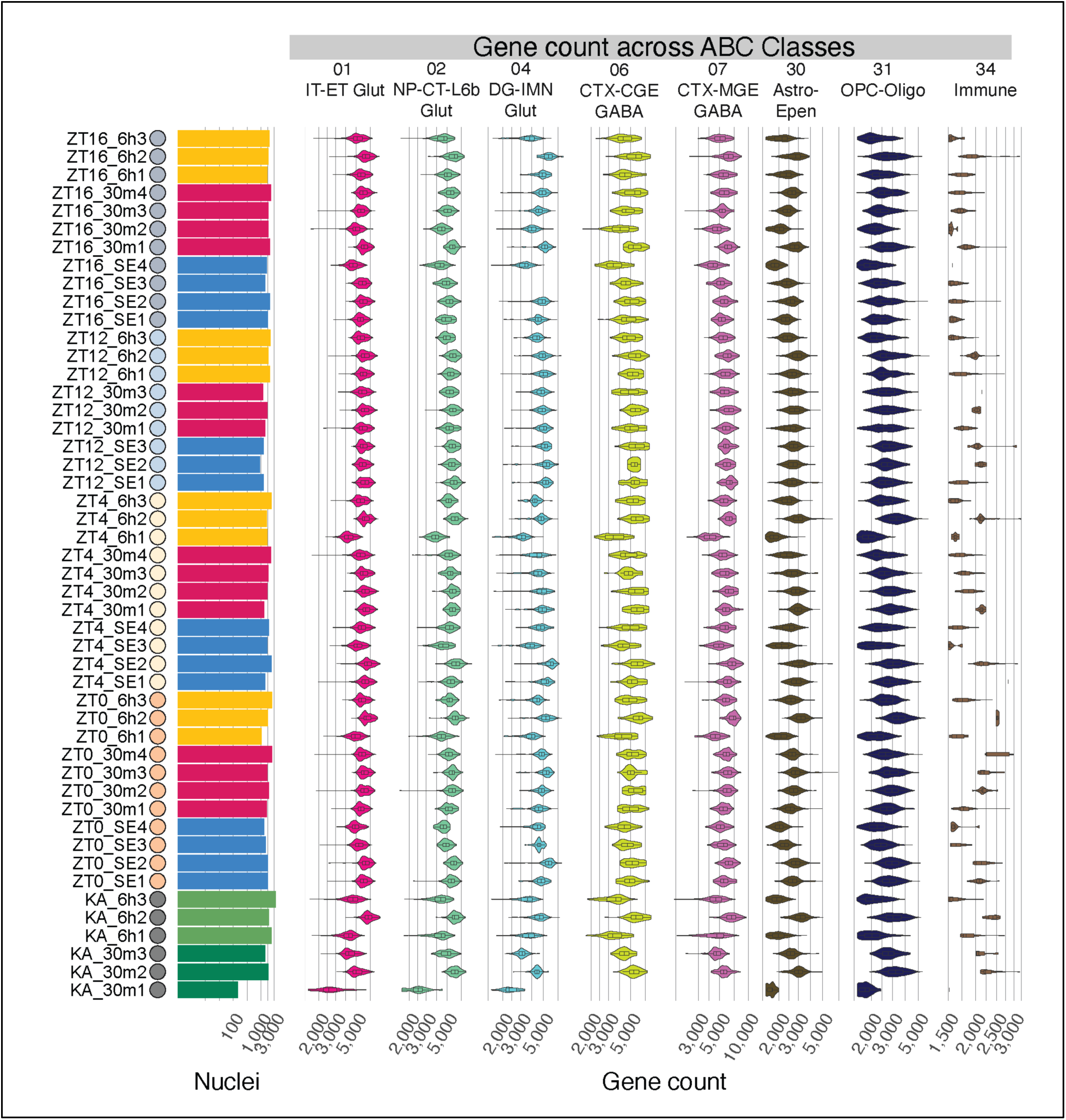
Gene counts among ABC cell Classes for each biological replicate.

**Figure 2 — figure supplement 1.**
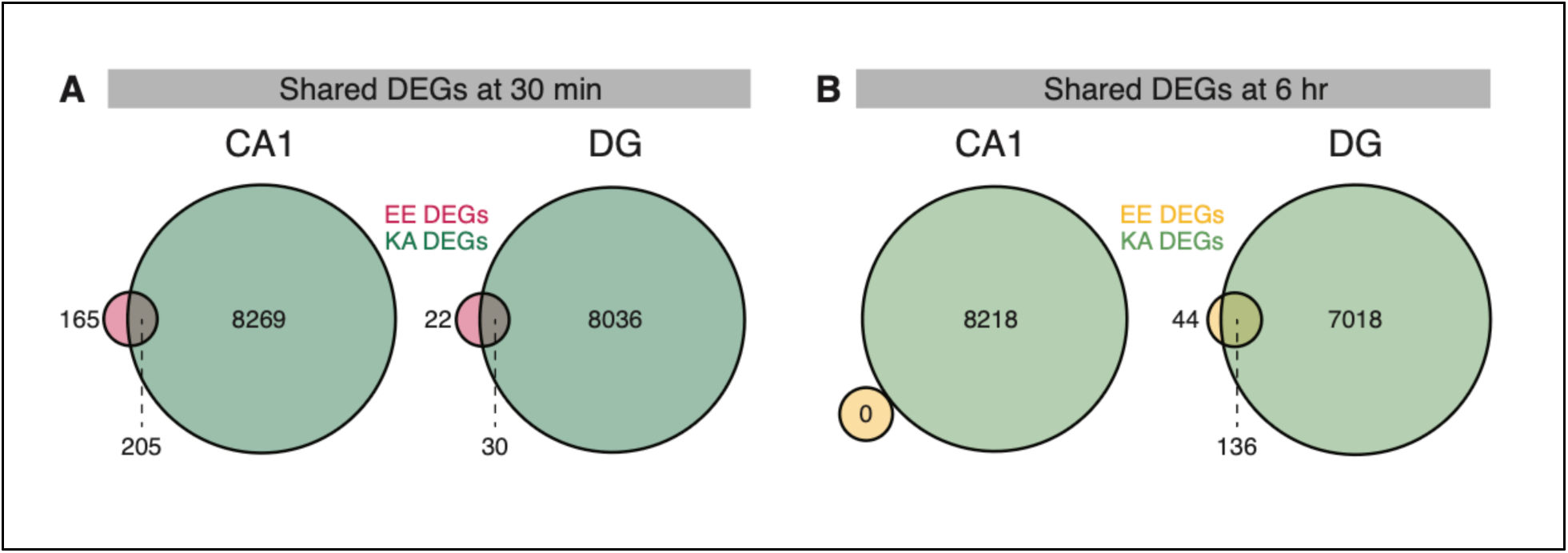
DGE analysis as in Figure 2A-B, but using limma- voom for pseudobulk DGE analysis instead of DESeq2.

**Figure 2 — figure supplement 2.**
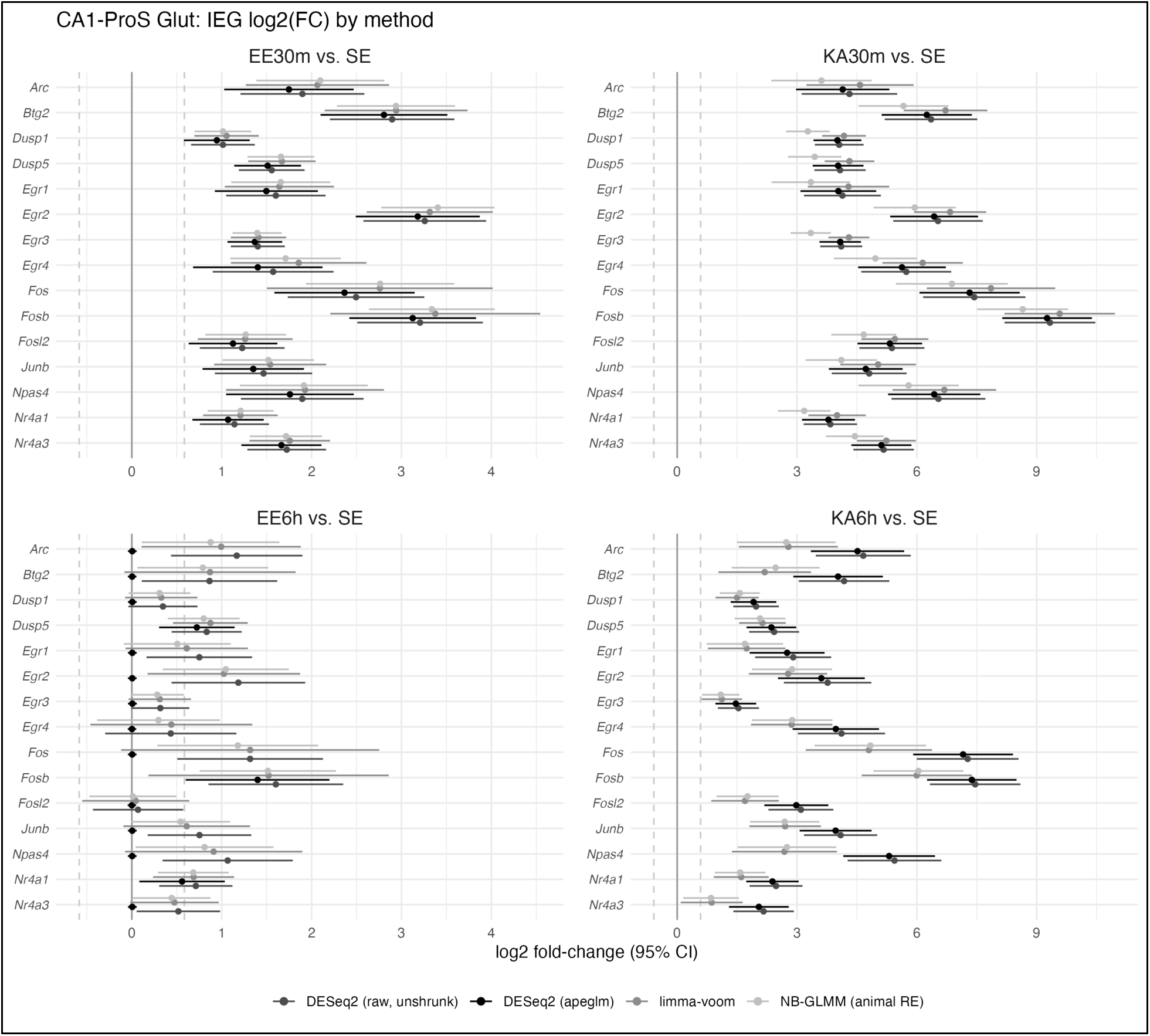
Estimations of IEG expression among different statistical frameworks within the CA1 population. (A) Estimated log2 fold-change in expression for IEGs in EE 30 min vs SE using DESeq2 unshrunken values (raw), DESeq2 shrunken values (apeglm), limma-voom, or a negative-binomial generalized linear mixed model (NB-GLMM). Dashed lines represent the fold-change threshold required to be considered a DEG (log2(FC) = 0.585, or 50% change in absolute expression). DESeq2 shrunken values were used for all analyses within main figures. (B-D) As in A, but for KA 30 min vs SE, EE 6h vs SE, and KA 6h vs SE.

**Figure 2 — figure supplement 3.**
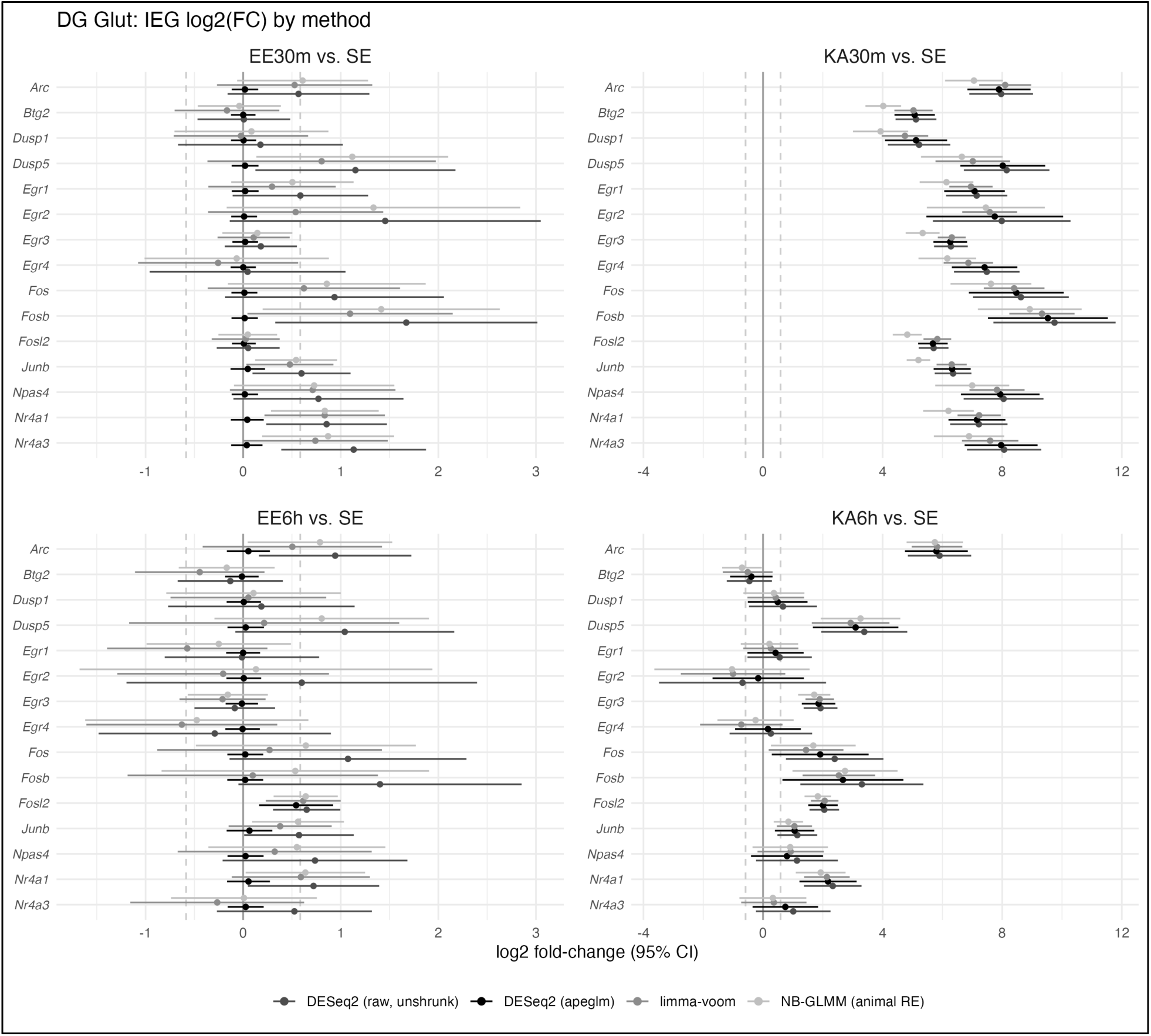
Estimations of IEG expression among different statistical frameworks within the DG population.

**Figure 2 — figure supplement 4.**
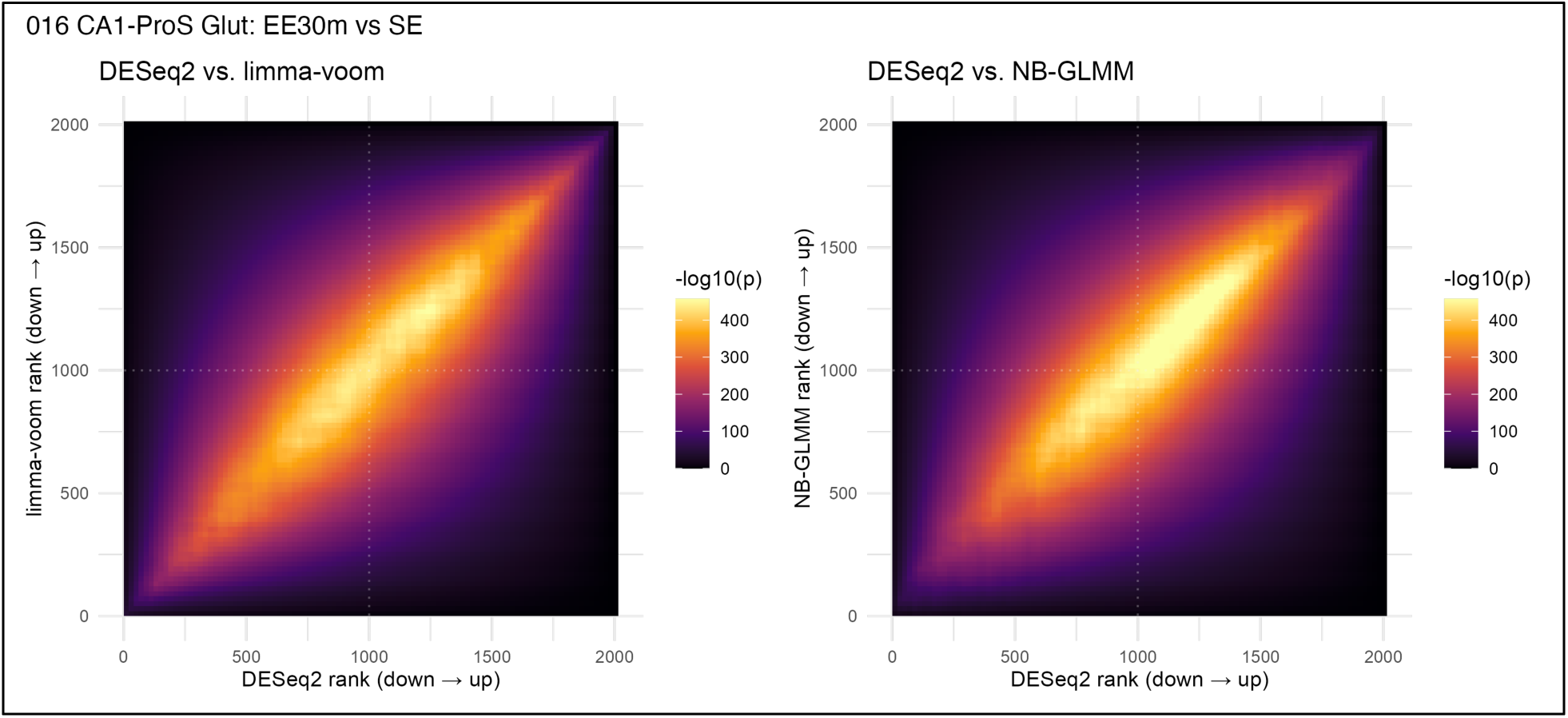
Comparing log2 fold-change estimations between DESeq2 vs limma-voom or NB-GLMM within CA1. (A) Rank-rank hypergeometric overlap (RRHO) plot comparing ranked gene vectors with expression values estimated by DESeq2 and limma-voom. Labeled vectors contained signed log2(FC) values x adjusted p-values. A Fisher’s test using the underlying hypergeometric distribution was used to determine the similarity of gene ranks between methods in bins of 100 genes along the vector, as in ref ^73^. Spearman’s *ρ* = 0.985. (B) As in A, but for DESeq2 vs NB-GLMM. Spearman’s *ρ* = 0.985.

**Figure 2 — figure supplement 5.**
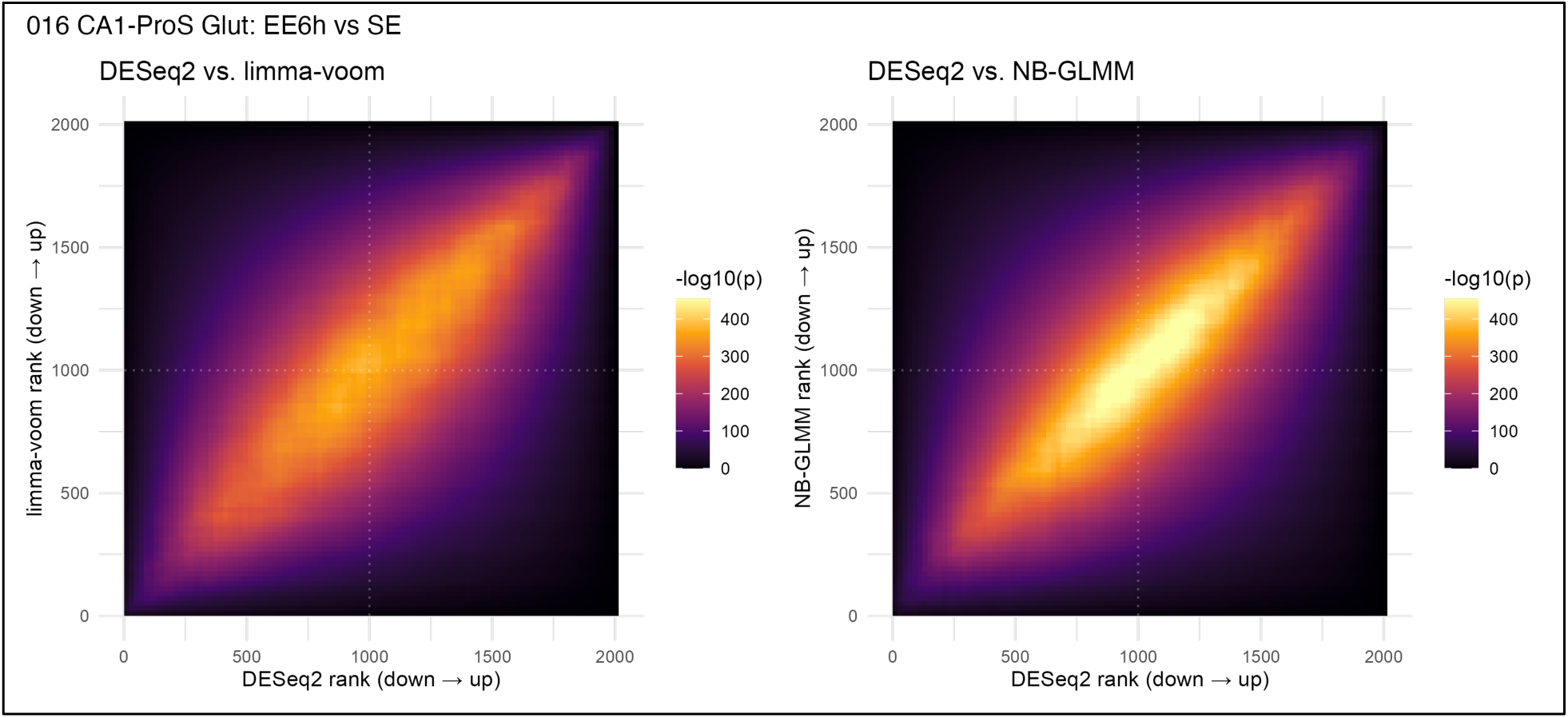
Comparing log2 fold-change estimations between DESeq2 vs limma-voom or NB-GLMM within DG. (A-B) As in **Figure 2 — figure supplement 5,** but for EE 6h. Spearman’s *ρ* = 0.964 in (A) and *ρ* = 0.983 in (B).

**Figure 2 — figure supplement 6.**
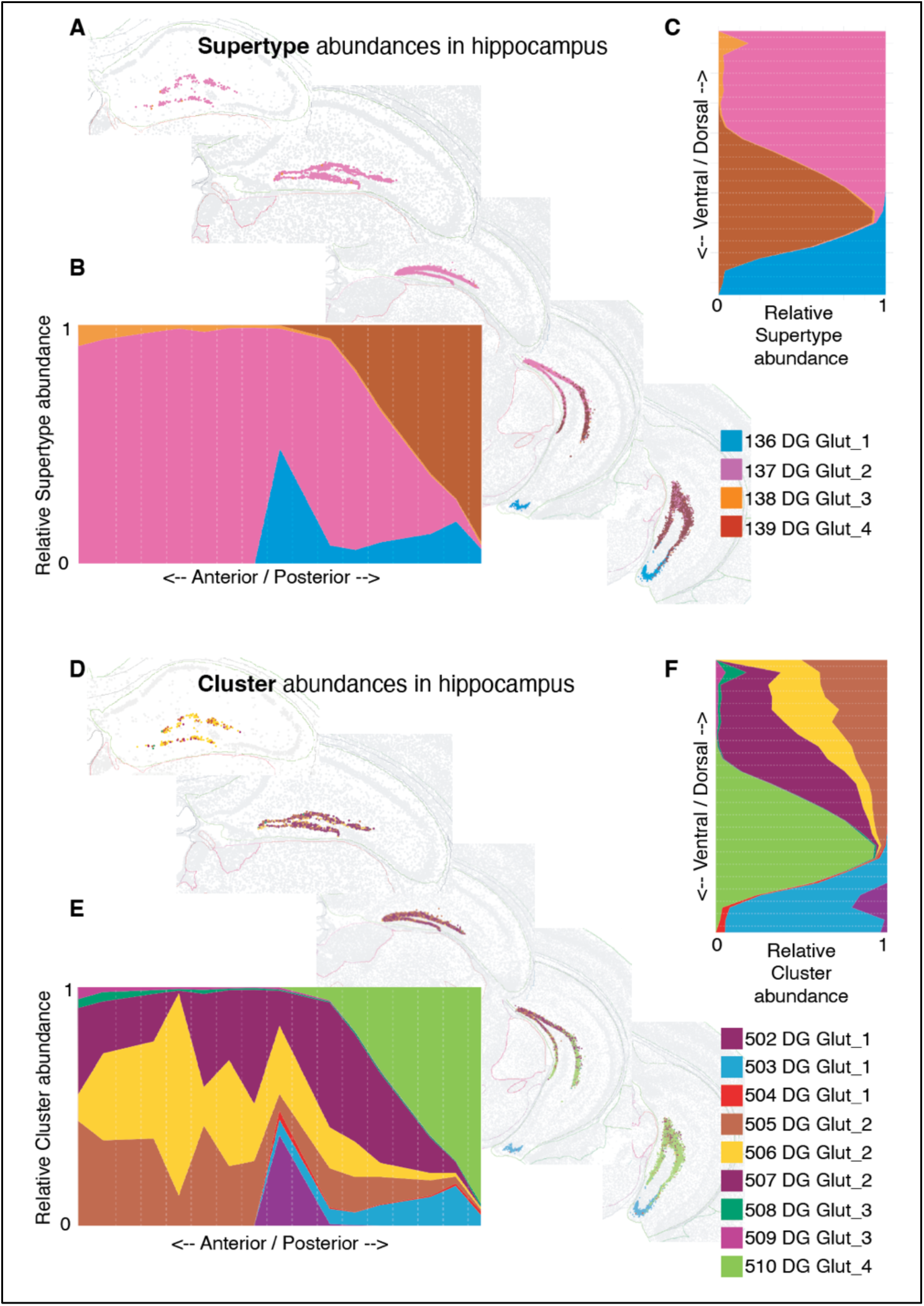
Spatial distribution of dentate granule cells within the Yao et al. 2023 ABC MERFISH atlas. (A-C) Supertype `138 DG Glut_3` (orange) is a rare population biased toward anterior/dorsal hippocampus. (D-F) Clusters `508` and `50G DG Glut_3` are sub-populations of 138 DG Glut_3 roughly co-equal in proportion and distribution.

**Figure 2 — figure supplement 7.**
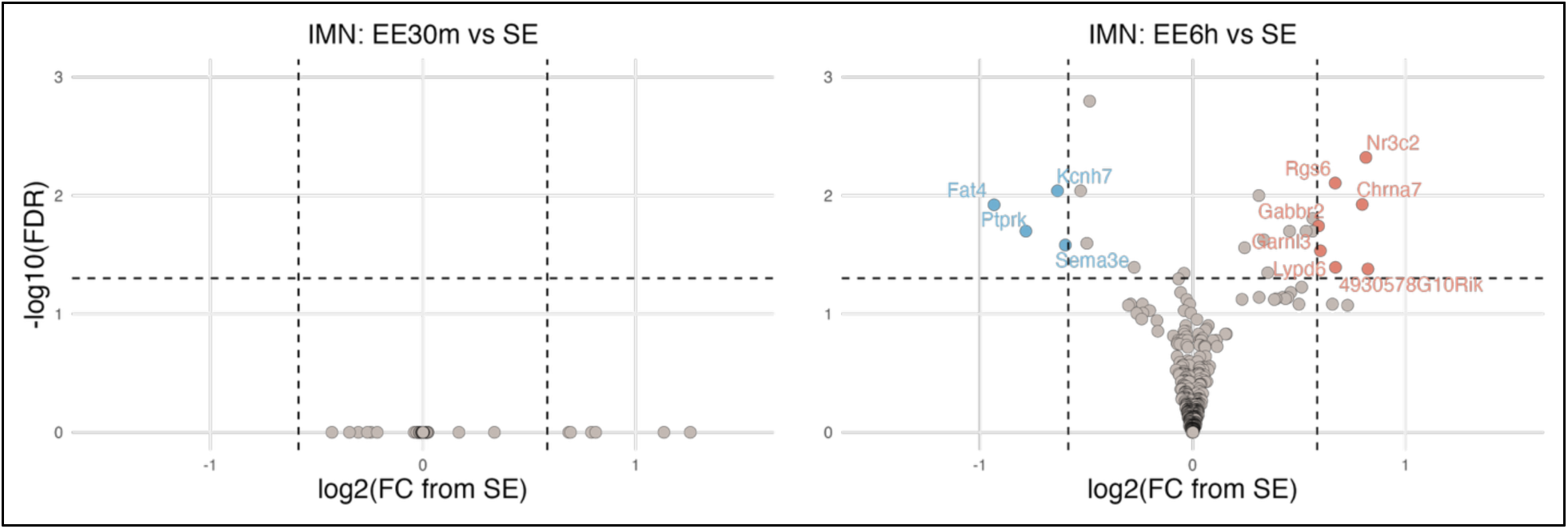
DEGs in immature dentate granule cells (IMN) after 30 minutes or 6 hours of enriched environment.

**Figure 3 — figure supplement 1.**
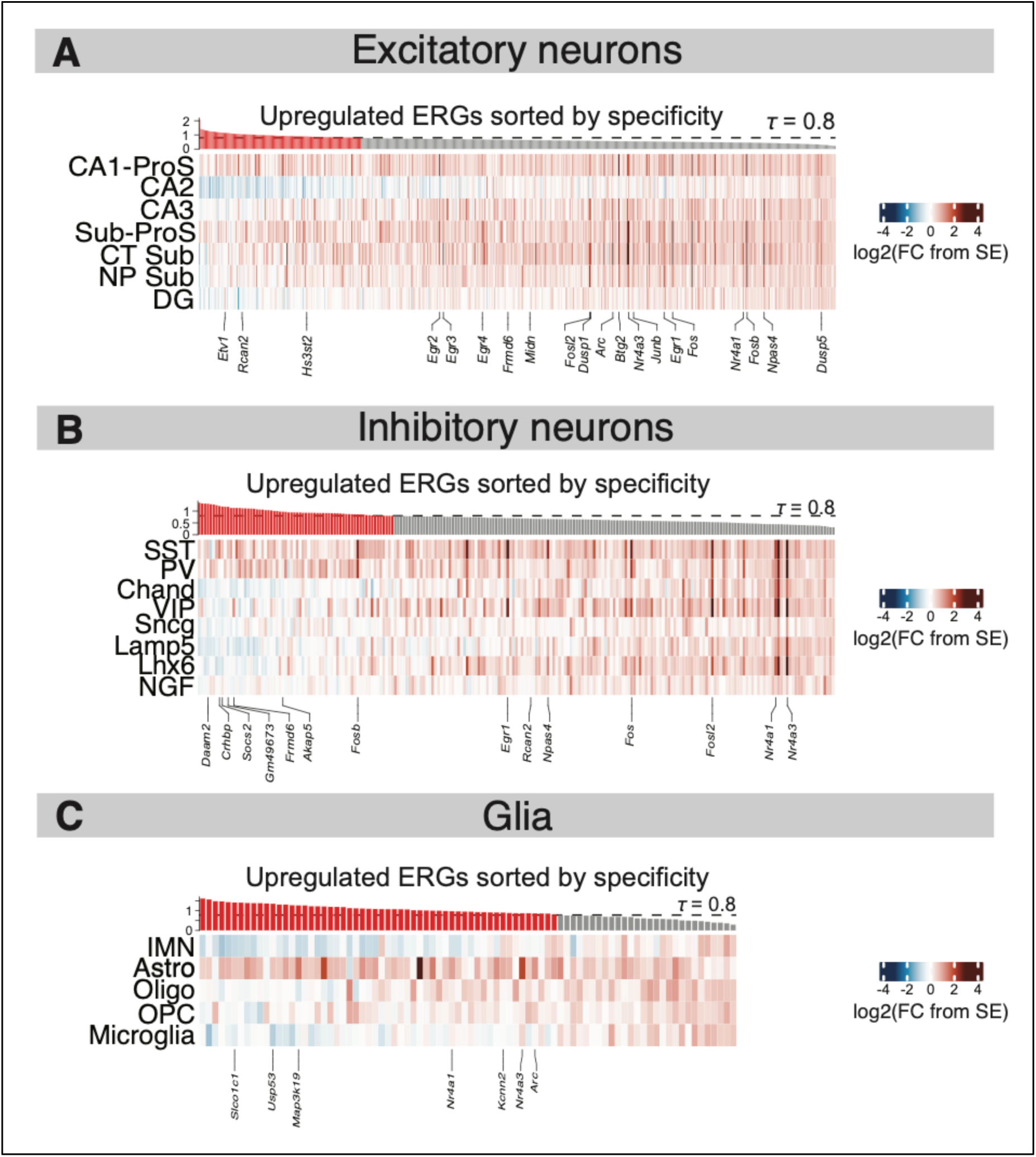
Tau heatmaps for upregulated ERGs ascertained using limma-voom. (A) As in Figure 3C, but instead using limma-voom instead of DESeq2 to detect DEGs. Log2(FC) values represent difference in mean log2-CPM in EE30m vs SE. (B-C) As in A, but for inhibitory neurons and glia.

**Figure 4 — figure supplement 1.**
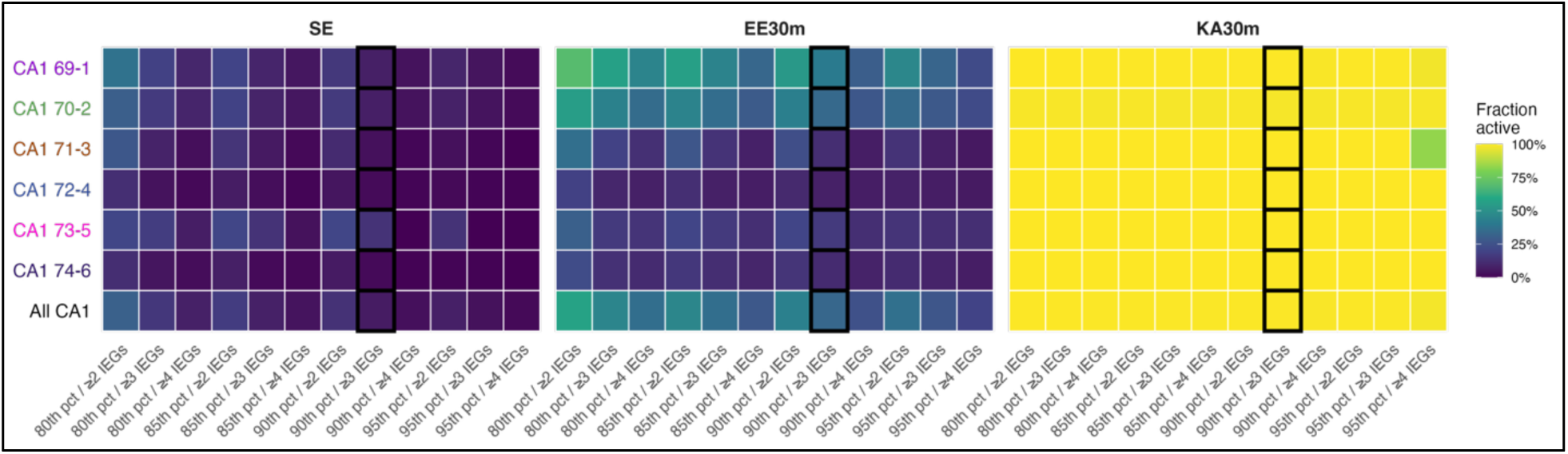
Robustness of active cell detection within CA1. The fraction of active CA1 cells when manipulating the number of genes or expression- percentile threshold required to classify a cell as active. Results are shown for each CA1 Supertype and total population. The black outlines denotes thresholds used in Figure 4B.

**Figure 4 — figure supplement 2.**
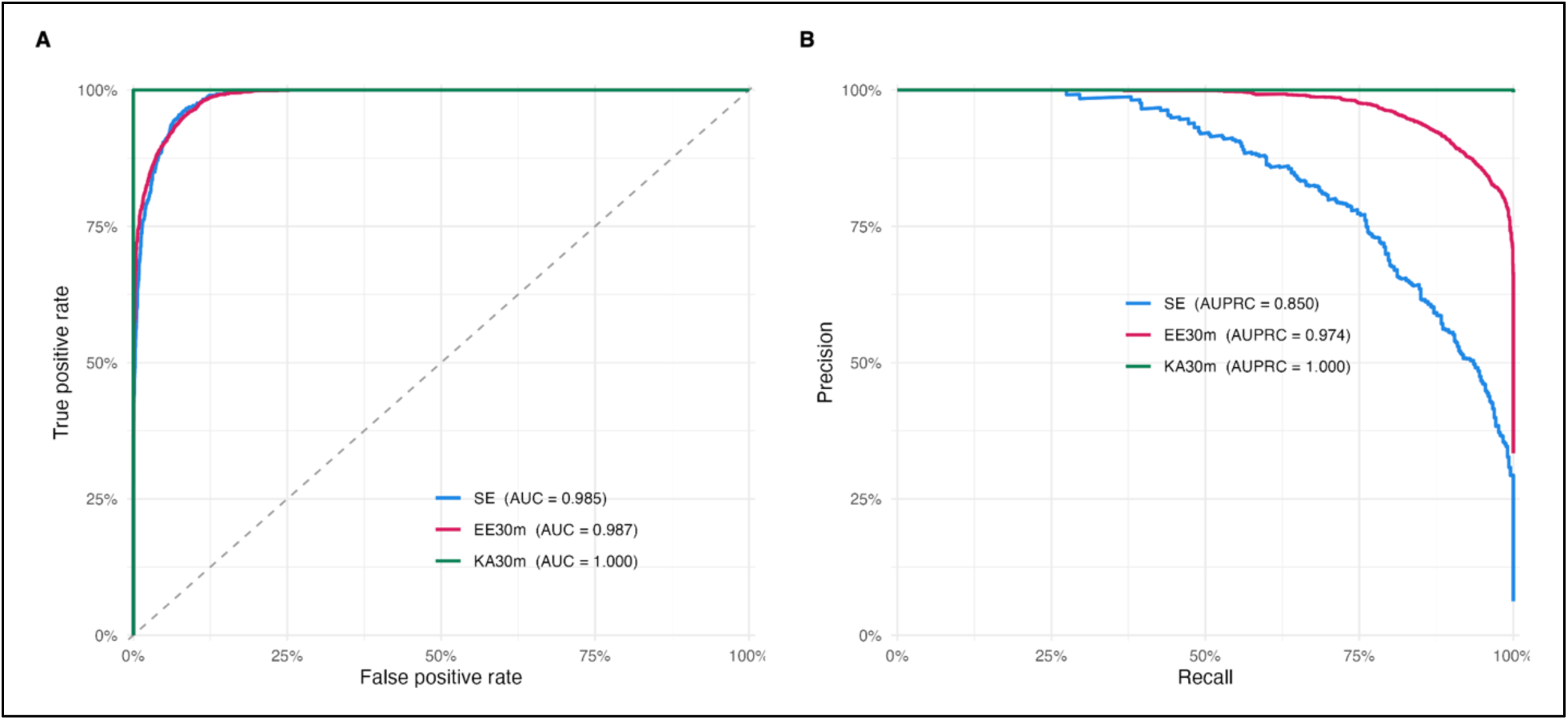
Classifying active cells using a continuous IEG score. Receiver operating characteristic (ROC) and precision-recall (PR) curves analyzing the accuracy of active vs inactive classification in CA1 cells using a continuous IEG score (**Methods**). Ground-truth classifications correspond to those used in Figure 4B.

**Figure 6 — figure supplement 1.**
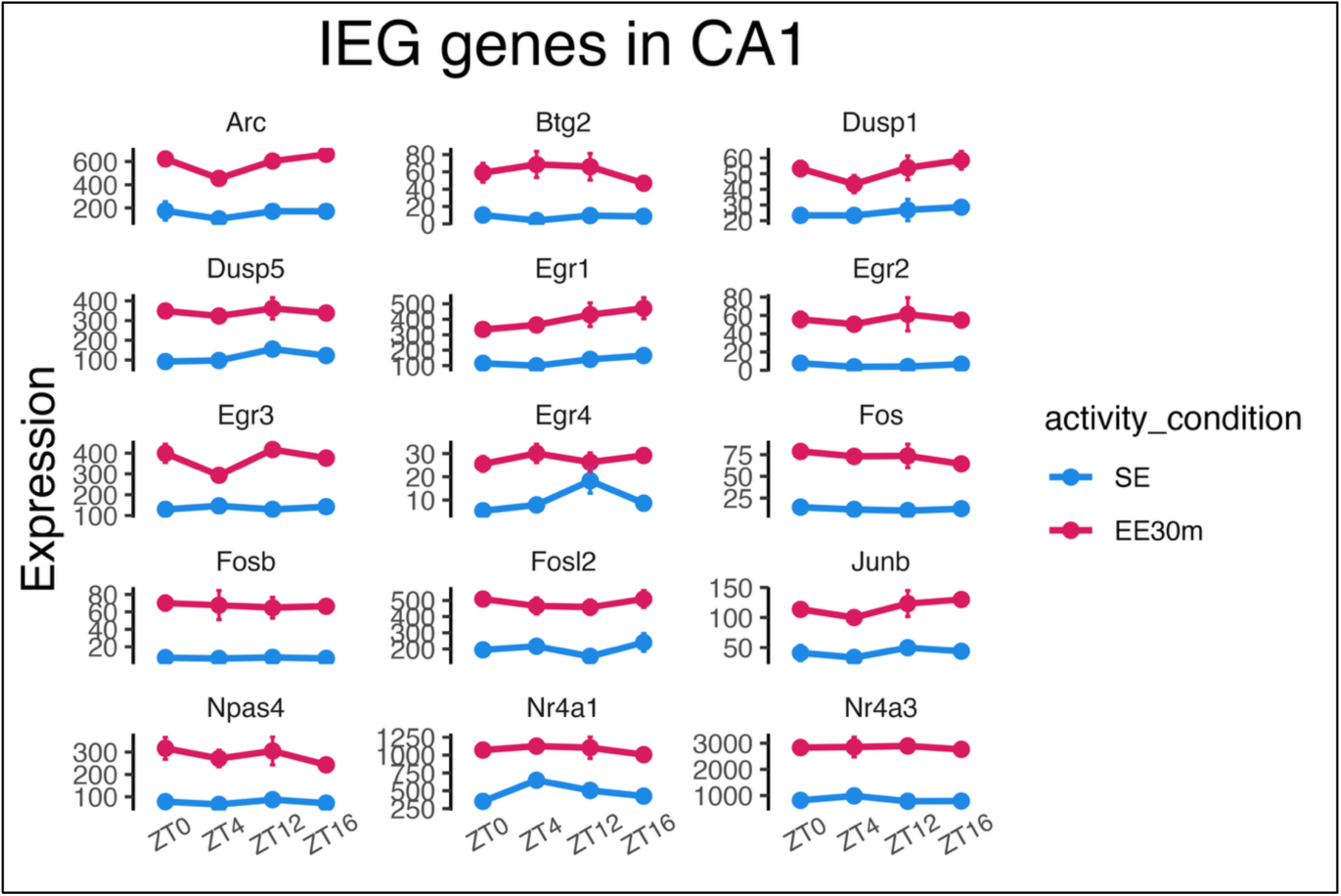
Pseudobulked expression of IEGs across the light- dark cycle within CA1 in SE and after EE 30 min.

**Figure 6 — figure supplement 2.**
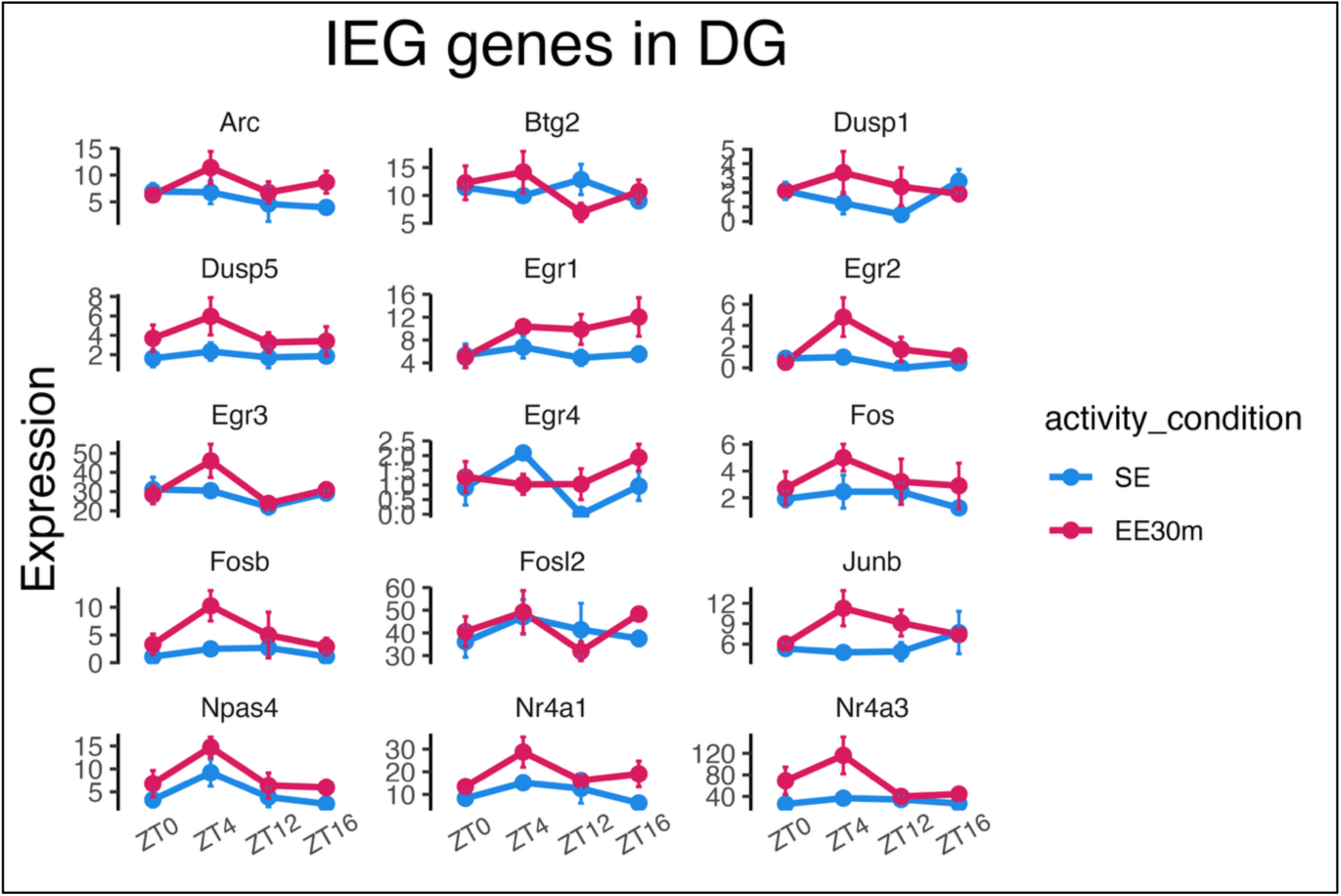
Pseudobulked expression of IEGs across the light-dark cycle within DG in SE and after EE 30 min.

## SUPPLEMENTAL ATTACHMENTS

**Supplemental item 1.**
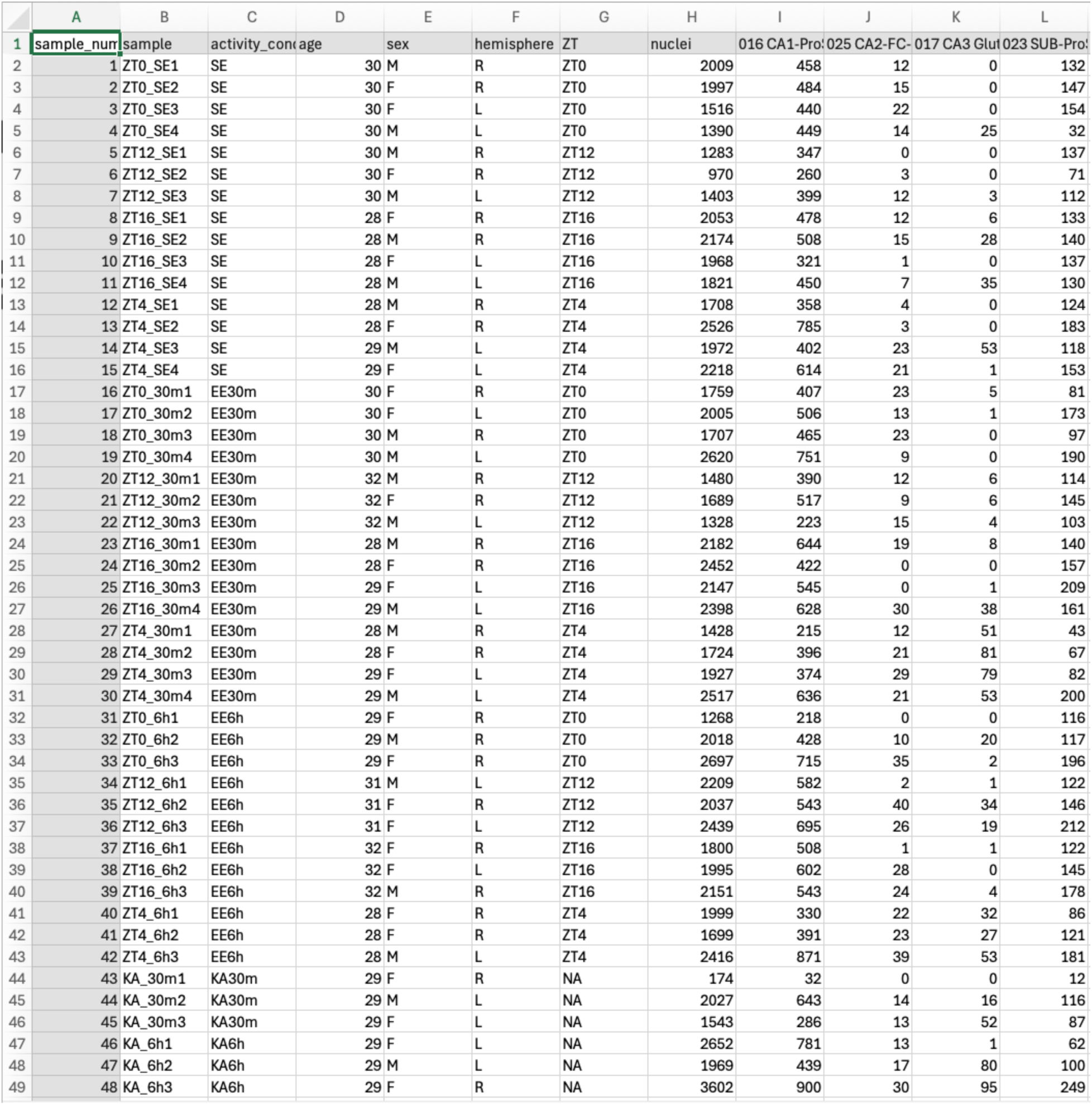
Biological replicates, metadata, and nuclei counts per subclass.

**Supplemental item 2.**
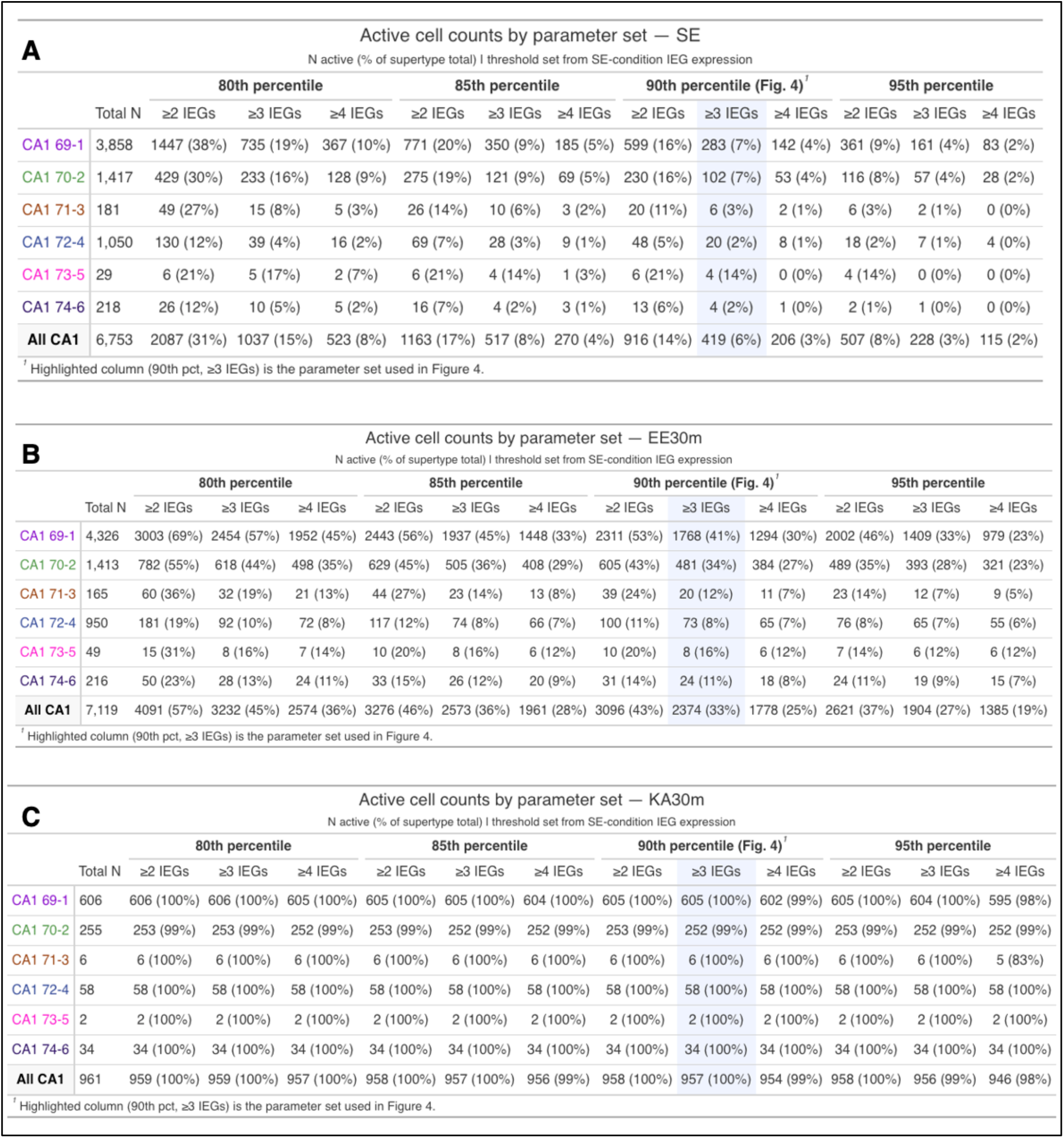
Parameter sweet values associated with Figure 4 — figure supplement 1.

## References

1. Greenberg, M. E. & Ziff, E. B. Stimulation of 3T3 cells induces transcription of the c-fos proto-oncogene. Nature 311, 433–438 (1984).

2. Tyssowski, K. M. et al. Different Neuronal Activity Patterns Induce Different Gene Expression Programs. Neuron 98, 530–546.e11 (2018).

3. Guzowski, J. F., McNaughton, B. L., Barnes, C. A. & Worley, P. F. Environment-specific expression of the immediate-early gene Arc in hippocampal neuronal ensembles. Nat Neurosci 2, 1120–1124 (1999).

4. Tullai, J. W. et al. Immediate-Early and Delayed Primary Response Genes Are Distinct in Function and Genomic Architecture*. Journal of Biological Chemistry 282, 23981– 23995 (2007).

5. Lituma, P. J., Singer, R. H., Das, S. & Castillo, P. E. Real-time imaging of Arc/Arg3.1 transcription ex vivo reveals input-specific immediate early gene dynamics. Proceedings of the National Academy of Sciences 119, e2123373119 (2022).

6. Akiki, R. M. et al. A long noncoding eRNA forms R-loops to shape emotional experience–induced behavioral adaptation. Science 386, 1282–1289 (2024).

7. Yao, Z. et al. A high-resolution transcriptomic and spatial atlas of cell types in the whole mouse brain. Nature 624, 317–332 (2023).

8. Langlieb, J. et al. The molecular cytoarchitecture of the adult mouse brain. NaturSingle-cell analysis of experience-dependent transcriptomic states in the mouse visual cortex. e 624, 333–342 (2023).

9. Lacar, B. et al. Nuclear RNA-seq of single neurons reveals molecular signatures of activation. Nat Commun 7, 11022 (2016).

10. Hrvatin, S. et al. Nat Neurosci 21, 120–129 (2018).

11. Jarrard, L. E. On the role of the hippocampus in learning and memory in the rat. Behavioral and Neural Biology 60, 9–26 (1993).

12. O’Keefe, J. & Dostrovsky, J. The hippocampus as a spatial map. Preliminary evidence from unit activity in the freely-moving rat. Brain Research 34, 171–175 (1971).

13. Nádasdy, Z., Hirase, H., Czurkó, A., Csicsvari, J. & Buzsáki, G. Replay and Time Compression of Recurring Spike Sequences in the Hippocampus. J. Neurosci. 19, 9497–9507 (1999).

14. Buzsáki, G. Hippocampal sharp wave-ripple: A cognitive biomarker for episodic memory and planning. Hippocampus 25, 1073–1188 (2015).

15. Joo, H. R. & Frank, L. M. The hippocampal sharp wave–ripple in memory retrieval for immediate use and consolidation. Nat Rev Neurosci 19, 744–757 (2018).

16. Carr, M. F., Jadhav, S. P. & Frank, L. M. Hippocampal replay in the awake state: a potential substrate for memory consolidation and retrieval. Nat Neurosci 14, 147– 153 (2011).

17. Pfeiffer, B. E. The content of hippocampal “replay”. Hippocampus 30, 6–18 (2020).

18. Bellfy, L. et al. The clock gene Per1 may exert diurnal control over hippocampal memory consolidation. Neuropsychopharmacol. 48, 1789–1797 (2023).

19. Kwapis, J. L. et al. Epigenetic regulation of the circadian gene Per1 contributes to age-related changes in hippocampal memory. Nat Commun 9, 3323 (2018).

20. Harvey, J. R. M., Plante, A. E. & Meredith, A. L. Ion Channels Controlling Circadian Rhythms in Suprachiasmatic Nucleus Excitability. Physiological Reviews 100, 1415– 1454 (2020).

21. Gonzalez, J. C. et al. Circadian regulation of dentate gyrus excitability mediated by G-protein signaling. Cell Reports 42, 112039 (2023).

22. Flourakis, M. et al. A Conserved Bicycle Model for Circadian Clock Control of Membrane Excitability. Cell 162, 836–848 (2015).

23. Birnie, M. T. et al. Circadian regulation of hippocampal function is disrupted with corticosteroid treatment. Proceedings of the National Academy of Sciences 120, e2211996120 (2023).

24. Gerstner, J. R. et al. BMAL1 controls the diurnal rhythm and set point for electrical seizure threshold in mice. Front. Syst. Neurosci. 8, (2014).

25. Karoly, P. J. et al. Circadian and circaseptan rhythms in human epilepsy: a retrospective cohort study. The Lancet Neurology 17, 977–985 (2018).

26. Rawashdeh, O. et al. PERIOD1 coordinates hippocampal rhythms and memory processing with daytime. Hippocampus 24, 712–723 (2014).

27. Olmstead, J. A transcriptomic atlas of activity-dependent gene expression across the circadian cycle. (UC San Diego, 2025).

28. Rosenberg, A. B. et al. Single-cell profiling of the developing mouse brain and spinal cord with split-pool barcoding. Science 360, 176–182 (2018).

29. Fernandez-Albert, J. et al. Immediate and deferred epigenomic signatures of in vivo neuronal activation in mouse hippocampus. Nat Neurosci 22, 1718–1730 (2019).

30. Traunmüller, L. et al. Novel environment exposure drives temporally defined and region-specific chromatin accessibility and gene expression changes in the hippocampus. Nat Commun 16, 7787 (2025).

31. Johnson, A., Varberg, Z., Benhardus, J., Maahs, A. & Schrater, P. The hippocampus and exploration: dynamically evolving behavior and neural representations. Front. Hum. Neurosci. 6, (2012).

32. Diamantaki, M., Frey, M., Berens, P., Preston-Ferrer, P. & Burgalossi, A. Sparse activity of identified dentate granule cells during spatial exploration. eLife 5, e20252 (2016).

33. Chavlis, S., Petrantonakis, P. C. & Poirazi, P. Dendrites of dentate gyrus granule cells contribute to pattern separation by controlling sparsity: DENDRITIC ROLE IN PATTERN SEPARATION. Hippocampus 27, 89–110 (2017).

34. Lee, J. Q. et al. Sparsity of Population Activity in the Hippocampus Is Task-Invariant Across the Trisynaptic Circuit and Dorsoventral Axis. Hippocampus 35, e23651 (2025).

35. Borzello, M. et al. Assessments of dentate gyrus function: discoveries and debates. Nat. Rev. Neurosci. 24, 502–517 (2023).

36. Yanai, I. et al. Genome-wide midrange transcription profiles reveal expression level relationships in human tissue specification. Bioinformatics 21, 650–659 (2005).

37. Pollina, E. A. et al. A NPAS4–NuA4 complex couples synaptic activity to DNA repair Nature 1–10 (2023) doi:10.1038/s41586-023-05711-7.

38. Yao, Z. et al. A transcriptomic and epigenomic cell atlas of the mouse primary motor cortex. Nature 598, 103–110 (2021).

39. van Strien, N. M., Cappaert, N. L. M. & Witter, M. P. The anatomy of memory: an interactive overview of the parahippocampal–hippocampal network. Nat Rev Neurosci 10, 272–282 (2009).

40. Rolotti, S. V., Blockus, H., Sparks, F. T., Priestley, J. B. & Losonczy, A. Reorganization of CA1 dendritic dynamics by hippocampal sharp-wave ripples during learning. Neuron 110, 977–991.e4 (2022).

41. Foster, D. j., Morris, R. g. m. & Dayan, P. A model of hippocampally dependent navigation, using the temporal difference learning rule. Hippocampus 10, 1–16 (2000).

42. Dragoi, G. & Tonegawa, S. Preplay of future place cell sequences by hippocampal cellular assemblies. Nature 469, 397–401 (2011).

43. Hor, C. N. et al. Sleep–wake-driven and circadian contributions to daily rhythms in gene expression and chromatin accessibility in the murine cortex. Proc. Natl. Acad. Sci. U.S.A. 116, 25773–25783 (2019).

44. Reick, M., Garcia, J. A., Dudley, C. & McKnight, S. L. NPAS2: An Analog of Clock Operative in the Mammalian Forebrain. Science 293, 506–509 (2001).

45. Wang, L. M. C. et al. Expression of the Circadian Clock Gene Period2 in the Hippocampus: Possible Implications for Synaptic Plasticity and Learned Behaviour. ASN Neuro 1, AN20090020 (2009).

46. Jilg, A. et al. Temporal dynamics of mouse hippocampal clock gene expression support memory processing. Hippocampus 20, 377–388 (2010).

47. Moriya, S., Tahara, Y., Sasaki, H., Ishigooka, J. & Shibata, S. Housing under abnormal light–dark cycles attenuates day/night expression rhythms of the clock genes Per1, Per2, and Bmal1 in the amygdala and hippocampus of mice. Neuroscience Research 99, 16–21 (2015).

48. Bering, T., Blancas-Velazquez, A. S. & Rath, M. F. Circadian Clock Genes Are Regulated by Rhythmic Corticosterone at Physiological Levels in the Rat Hippocampus. Neuroendocrinology 113, 1076–1090 (2023).

49. Chiou, Y.-Y. et al. Mammalian Period represses and de-represses transcription by displacing CLOCK–BMAL1 from promoters in a Cryptochrome-dependent manner. Proc. Natl. Acad. Sci. U.S.A. 113, (2016).

50. Eckel-Mahan, K. L. et al. Circadian oscillation of hippocampal MAPK activity and cAMP: implications for memory persistence. Nat Neurosci 11, 1074–1082 (2008).

51. Cirelli, C. & Tononi, G. Gene expression in the brain across the sleep–waking cycle1. Brain Research 885, 303–321 (2000).

52. Phan, T. H., Chan, G. C.-K., Sindreu, C. B., Eckel-Mahan, K. L. & Storm, D. R. The Diurnal Oscillation of MAP (Mitogen-Activated Protein) Kinase and Adenylyl Cyclase Activities in the Hippocampus Depends on the Suprachiasmatic Nucleus. J. Neurosci. 31, 10640–10647 (2011).

53. Squair, J. W. et al. Confronting false discoveries in single-cell differential expression. Nat Commun 12, 5692 (2021).

54. Zimmerman, K. D., Espeland, M. A. & Langefeld, C. D. A practical solution to pseudoreplication bias in single-cell studies. Nat Commun 12, 738 (2021).

55. Denny, C. A. et al. Hippocampal Memory Traces Are Differentially Modulated by Experience, Time, and Adult Neurogenesis. Neuron 83, 189–201 (2014).

56. Sun, X. et al. Functionally Distinct Neuronal Ensembles within the Memory Engram. Cell 181, 410–423.e17 (2020).

57. Guenthner, C. J., Miyamichi, K., Yang, H. H., Heller, H. C. & Luo, L. Permanent Genetic Access to Transiently Active Neurons via TRAP: Targeted Recombination in Active Populations. Neuron 78, 773–784 (2013).

58. Sørensen, A. T. et al. A robust activity marking system for exploring active neuronal ensembles. eLife 5, e13918 (2016).

59. Patton, A. P., Hastings, M. H. & Smyllie, N. J. Cells and Circuits of the Suprachiasmatic Nucleus and the Control of Circadian Behaviour and Sleep. in Sleep and Clocks in Aging and Longevity (ed. Jagota, A.) 33–70 (Springer International Publishing, Cham, 2023). doi:10.1007/978-3-031-22468-3_2.

60. Patton, A. P. & Hastings, M. H. The suprachiasmatic nucleus. Current Biology 28, R816–R822 (2018).

61. Riggle, J. P. et al. Spontaneous Recovery of Circadian Organization in Mice Lacking a Core Component of the Molecular Clockwork. J Biol Rhythms 37, 94–109 (2022).

62. Tunnacliffe, E. & Chubb, J. R. What Is a Transcriptional Burst? Trends in Genetics 36, 288–297 (2020).

63. Corrigan, A. M. & Chubb, J. R. Regulation of Transcriptional Bursting by a Naturally Oscillating Signal. Current Biology 24, 205–211 (2014).

64. Coulon, A., Chow, C. C., Singer, R. H. & Larson, D. R. Eukaryotic transcriptional dynamics: from single molecules to cell populations. Nat Rev Genet 14, 10.1038/nrg3484 (2013).

65. Chen, L.-F. et al. Enhancer Histone Acetylation Modulates Transcriptional Bursting Dynamics of Neuronal Activity-Inducible Genes. Cell Reports 26, 1174–1188.e5 (2019).

66. Luo, J., Phan, T. X., Yang, Y., Garelick, M. G. & Storm, D. R. Increases in cAMP, MAPK Activity, and CREB Phosphorylation during REM Sleep: Implications for REM Sleep and Memory Consolidation. J. Neurosci. 33, 6460–6468 (2013).

67. Saraf, A., Luo, J., Morris, D. R. & Storm, D. R. Phosphorylation of Eukaryotic Translation Initiation Factor 4E and Eukaryotic Translation Initiation Factor 4E- binding Protein (4EBP) and Their Upstream Signaling Components Undergo Diurnal Oscillation in the Mouse Hippocampus. Journal of Biological Chemistry 289, 20129– 20138 (2014).

68. Rönnbäck, A., Dahlqvist, P., Bergström, S.-A. & Olsson, T. Diurnal effects of enriched environment on immediate early gene expression in the rat brain. Brain Research 1046, 137–144 (2005).

69. Bosco, D. B. et al. RNAseq analysis of hippocampal microglia after kainic acid-induced seizures. Mol Brain 11, 34 (2018).

70. Racine, R. J. Modification of seizure activity by electrical stimulation: II. Motor seizure. Electroencephalography and Clinical Neurophysiology 32, 281–294 (1972).

71. McGinnis, C. S., Murrow, L. M. & Gartner, Z. J. DoubletFinder: Doublet Detection in Single-Cell RNA Sequencing Data Using Artificial Nearest Neighbors. Cell Systems 8, 329–337.e4 (2019).

72. Love, M. I., Huber, W. & Anders, S. Moderated estimation of fold change and dispersion for RNA-seq data with DESeq2. Genome Biology 15, 550 (2014).

73. Piron, A. et al. RedRibbon: A new rank–rank hypergeometric overlap for gene and transcript expression signatures. Life Science Alliance 7, (2024).

74. Gu, Z., Eils, R. & Schlesner, M. Complex heatmaps reveal patterns and correlations in multidimensional genomic data. Bioinformatics 32, 2847–2849 (2016).

75. Robin, X. et al. pROC: an open-source package for R and S+ to analyze and compare ROC curves. BMC Bioinformatics 12, 77 (2011).

76. Grau, J., Grosse, I. & Keilwagen, J. PRROC: computing and visualizing precision- recall and receiver operating characteristic curves in R. Bioinformatics 31, 2595– 2597 (2015).

