## Supplemental Item 2 for "Intersection of transient cell states with stable cell types in hippocampus"

**A**

### Active cell counts by parameter set — SE

N active (% of supertype total) | threshold set from SE-condition IEG expression

| | | 80th percentile | | | 85th percentile | | | 90th percentile (Fig. 4)[1] | | | 95th percentile | | |
|---|---|---|---|---|---|---|---|---|---|---|---|---|---|
| | Total N | ≥2 IEGs | ≥3 IEGs | ≥4 IEGs | ≥2 IEGs | ≥3 IEGs | ≥4 IEGs | ≥2 IEGs | ≥3 IEGs | ≥4 IEGs | ≥2 IEGs | ≥3 IEGs | ≥4 IEGs |
| CA1 69-1 | 3,858 | 1447 (38%) | 735 (19%) | 367 (10%) | 771 (20%) | 350 (9%) | 185 (5%) | 599 (16%) | 283 (7%) | 142 (4%) | 361 (9%) | 161 (4%) | 83 (2%) |
| CA1 70-2 | 1,417 | 429 (30%) | 233 (16%) | 128 (9%) | 275 (19%) | 121 (9%) | 69 (5%) | 230 (16%) | 102 (7%) | 53 (4%) | 116 (8%) | 57 (4%) | 28 (2%) |
| CA1 71-3 | 181 | 49 (27%) | 15 (8%) | 5 (3%) | 26 (14%) | 10 (6%) | 3 (2%) | 20 (11%) | 6 (3%) | 2 (1%) | 6 (3%) | 2 (1%) | 0 (0%) |
| CA1 72-4 | 1,050 | 130 (12%) | 39 (4%) | 16 (2%) | 69 (7%) | 28 (3%) | 9 (1%) | 48 (5%) | 20 (2%) | 8 (1%) | 18 (2%) | 7 (1%) | 4 (0%) |
| CA1 73-5 | 29 | 6 (21%) | 5 (17%) | 2 (7%) | 6 (21%) | 4 (14%) | 1 (3%) | 6 (21%) | 4 (14%) | 0 (0%) | 4 (14%) | 0 (0%) | 0 (0%) |
| CA1 74-6 | 218 | 26 (12%) | 10 (5%) | 5 (2%) | 16 (7%) | 4 (2%) | 3 (1%) | 13 (6%) | 4 (2%) | 1 (0%) | 2 (1%) | 1 (0%) | 0 (0%) |
| **All CA1** | 6,753 | 2087 (31%) | 1037 (15%) | 523 (8%) | 1163 (17%) | 517 (8%) | 270 (4%) | 916 (14%) | 419 (6%) | 206 (3%) | 507 (8%) | 228 (3%) | 115 (2%) |

[1] Highlighted column (90th pct, ≥3 IEGs) is the parameter set used in Figure 4.

**B**

### Active cell counts by parameter set — EE30m

N active (% of supertype total) | threshold set from SE-condition IEG expression

| | | 80th percentile | | | 85th percentile | | | 90th percentile (Fig. 4)[1] | | | 95th percentile | | |
|---|---|---|---|---|---|---|---|---|---|---|---|---|---|
| | Total N | ≥2 IEGs | ≥3 IEGs | ≥4 IEGs | ≥2 IEGs | ≥3 IEGs | ≥4 IEGs | ≥2 IEGs | ≥3 IEGs | ≥4 IEGs | ≥2 IEGs | ≥3 IEGs | ≥4 IEGs |
| CA1 69-1 | 4,326 | 3003 (69%) | 2454 (57%) | 1952 (45%) | 2443 (56%) | 1937 (45%) | 1448 (33%) | 2311 (53%) | 1768 (41%) | 1294 (30%) | 2002 (46%) | 1409 (33%) | 979 (23%) |
| CA1 70-2 | 1,413 | 782 (55%) | 618 (44%) | 498 (35%) | 629 (45%) | 505 (36%) | 408 (29%) | 605 (43%) | 481 (34%) | 384 (27%) | 489 (35%) | 393 (28%) | 321 (23%) |
| CA1 71-3 | 165 | 60 (36%) | 32 (19%) | 21 (13%) | 44 (27%) | 23 (14%) | 13 (8%) | 39 (24%) | 20 (12%) | 11 (7%) | 23 (14%) | 12 (7%) | 9 (5%) |
| CA1 72-4 | 950 | 181 (19%) | 92 (10%) | 72 (8%) | 117 (12%) | 74 (8%) | 66 (7%) | 100 (11%) | 73 (8%) | 65 (7%) | 76 (8%) | 65 (7%) | 55 (6%) |
| CA1 73-5 | 49 | 15 (31%) | 8 (16%) | 7 (14%) | 10 (20%) | 8 (16%) | 6 (12%) | 10 (20%) | 8 (16%) | 6 (12%) | 7 (14%) | 6 (12%) | 6 (12%) |
| CA1 74-6 | 216 | 50 (23%) | 28 (13%) | 24 (11%) | 33 (15%) | 26 (12%) | 20 (9%) | 31 (14%) | 24 (11%) | 18 (8%) | 24 (11%) | 19 (9%) | 15 (7%) |
| **All CA1** | 7,119 | 4091 (57%) | 3232 (45%) | 2574 (36%) | 3276 (46%) | 2573 (36%) | 1961 (28%) | 3096 (43%) | 2374 (33%) | 1778 (25%) | 2621 (37%) | 1904 (27%) | 1385 (19%) |

[1] Highlighted column (90th pct, ≥3 IEGs) is the parameter set used in Figure 4.

**C**

### Active cell counts by parameter set — KA30m

N active (% of supertype total) | threshold set from SE-condition IEG expression

| | | 80th percentile | | | 85th percentile | | | 90th percentile (Fig. 4)[1] | | | 95th percentile | | |
|---|---|---|---|---|---|---|---|---|---|---|---|---|---|
| | Total N | ≥2 IEGs | ≥3 IEGs | ≥4 IEGs | ≥2 IEGs | ≥3 IEGs | ≥4 IEGs | ≥2 IEGs | ≥3 IEGs | ≥4 IEGs | ≥2 IEGs | ≥3 IEGs | ≥4 IEGs |
| CA1 69-1 | 606 | 606 (100%) | 606 (100%) | 605 (100%) | 605 (100%) | 605 (100%) | 604 (100%) | 605 (100%) | 605 (100%) | 602 (99%) | 605 (100%) | 604 (100%) | 595 (98%) |
| CA1 70-2 | 255 | 253 (99%) | 253 (99%) | 252 (99%) | 253 (99%) | 252 (99%) | 252 (99%) | 253 (99%) | 252 (99%) | 252 (99%) | 253 (99%) | 252 (99%) | 252 (99%) |
| CA1 71-3 | 6 | 6 (100%) | 6 (100%) | 6 (100%) | 6 (100%) | 6 (100%) | 6 (100%) | 6 (100%) | 6 (100%) | 6 (100%) | 6 (100%) | 6 (100%) | 5 (83%) |
| CA1 72-4 | 58 | 58 (100%) | 58 (100%) | 58 (100%) | 58 (100%) | 58 (100%) | 58 (100%) | 58 (100%) | 58 (100%) | 58 (100%) | 58 (100%) | 58 (100%) | 58 (100%) |
| CA1 73-5 | 2 | 2 (100%) | 2 (100%) | 2 (100%) | 2 (100%) | 2 (100%) | 2 (100%) | 2 (100%) | 2 (100%) | 2 (100%) | 2 (100%) | 2 (100%) | 2 (100%) |
| CA1 74-6 | 34 | 34 (100%) | 34 (100%) | 34 (100%) | 34 (100%) | 34 (100%) | 34 (100%) | 34 (100%) | 34 (100%) | 34 (100%) | 34 (100%) | 34 (100%) | 34 (100%) |
| **All CA1** | 961 | 959 (100%) | 959 (100%) | 957 (100%) | 958 (100%) | 957 (100%) | 956 (99%) | 958 (100%) | 957 (100%) | 954 (99%) | 958 (100%) | 956 (99%) | 946 (98%) |

[1] Highlighted column (90th pct, ≥3 IEGs) is the parameter set used in Figure 4.
